# The ER protein translocation channel subunit Sbh1 controls virulence of *Cryptococcus neoformans*

**DOI:** 10.1101/2022.06.01.494298

**Authors:** Felipe H. Santiago-Tirado, Thomas Hurtaux, Jennifer Geddes-McAlister, Duy Nguyen, Volkhard Helms, Tamara L. Doering, Karin Römisch

**Author notes:** Department of Biological Sciences, University of Notre Dame, Notre Dame, IN, USA.

## Abstract

The fungal pathogen *Cryptococcus neoformans* is distinguished by a cell wall-anchored polysaccharide capsule that is critical for virulence. Biogenesis of both cell wall and capsule relies on the secretory pathway. Protein secretion begins with polypeptide translocation across the endoplasmic reticulum (ER) membrane through a highly conserved channel formed by three proteins: Sec61, Sbh1, and Sss1. Sbh1, the most divergent, contains multiple phosphorylation sites, which may allow it to regulate entry into the secretory pathway in a species- and protein-specific manner. Absence of *SBH1* causes a cell-wall defect in both *Saccharomyces cerevisiae* and *C. neoformans*, although other phenotypes differ. Notably, proteomic analysis showed that when cryptococci are grown in conditions that mimic aspects of the mammalian host environment (tissue culture medium, 37 °C, 5% CO_2_), a set of secretory and transmembrane proteins is upregulated in wild-type, but not in *Δsbh1* mutant cells. The Sbh1-dependent proteins show specific features of their ER targeting sequences that likely cause them to transit less efficiently into the secretory pathway. Many also act in cell-wall biogenesis, while several are known virulence factors; consistent with these observations, the *C. neoformans Δsbh1* mutant is avirulent in a mouse infection model. We conclude that, in the context of conditions encountered during infection, Sbh1 controls the entry of virulence factors into the secretory pathway of *C. neoformans*, and thereby regulates fungal pathogenicity.

**Importance:** *Cryptococcus neoformans* is a yeast that causes almost 200,000 deaths worldwide each year, mainly of immunocompromised individuals. The surface structures of this pathogen, a protective cell wall surrounded by a polysaccharide capsule, are made and maintained by proteins that are synthesized inside the cell and travel outwards through the secretory pathway. A protein called Sbh1 is part of the machinery that determines which polypeptides enter this export pathway. We found that when Sbh1 is absent, both *C. neoformans* and the model yeast *S. cerevisiae* show cell wall defects. Lack of Sbh1 also changes the pattern of secretion of both transmembrane and soluble proteins, in a manner that depends on characteristics of their sequences. Notably, multiple proteins that are normally upregulated in conditions similar to those encountered during infection, including several needed for cryptococcal virulence, are no longer increased. Sbh1 thereby regulates the ability of this important pathogen to cause disease.

## INTRODUCTION

*Cryptococcus neoformans* is a haploid budding yeast that is ubiquitous in the environment, so that spores or dessicated cells are frequently inhaled (1, 2). In healthy individuals these infectious particles are generally cleared or establish an asymptomatic latent infection (3). If, however, the host is or becomes severely immunocompromised, the pathogen may grow and disseminate, both within and outside of host cells (4). Dissemination to the central nervous system results in a devastating meningitis, which causes almost 200,000 deaths each year worldwide (5).

Like other fungi, *C. neoformans* is protected by a multilayer cell wall, which is composed of a meshwork of chitin, chitosan, alpha and beta glucans, and mannoproteins (6). This flexible and dynamic structure responds to and protects the cell from environmental stresses, while accommodating morphogenesis. An additional role of the cryptococcal wall is to anchor an elaborate polysaccharide capsule that surrounds the cell (7, 8). This structure, which is unique among fungal pathogens, is required for cryptococcal virulence (1). Capsule polysaccharides are also shed from the cell into the surrounding environment, where they act to perturb the host immune response (9, 10).

The biogenesis of both cell wall and capsule in *C. neoformans* relies on the secretory pathway, which begins with protein translocation across the ER membrane through the highly conserved Sec61 channel (11). An N-terminal hydrophobic signal peptide or a transmembrane domain targets proteins to this channel, which is formed by three proteins: Sec61, Sbh1, and Sss1 in yeast (Fig. 1A) and Sec61*α*, Sec61ß, and Sec61*γ* in mammals (11, 12). *SEC61* and *SSS1* are essential in *S. cerevisiae* (13, 14). The 10 transmembrane helices of Sec61 form a protein-conducting channel in the ER membrane, stabilized by Sss1 which clamps around the Sec61 helix-bundle (Fig. 1A) (12, 15, 16). *SBH1* and its paralog *SBH2* are not essential, but deletion of both genes leads to temperature-sensitivity at 37°C in *S. cerevisiae* (17, 18). Sbh1/Sec61ß is a tail-anchored protein associated peripherally with the Sec61 channel (19) (Fig. 1A). Its conserved transmembrane domain is sufficient to rescue the double mutant growth defect at 37°C, while the role of its divergent cytosolic domain containing multiple phosphorylation sites is poorly understood (18, 20, 21). As is common for proteins with intrinsically unstructured domains, Sbh1/Sec61ß has multiple interactors: it binds to ribosomes and mediates association of the Sec61 channel with signal peptidase and signal recognition particle receptor (22, 23, 24, 25). In mammalian cells, the Sbh1 homolog Sec61ß also interacts with the Stimulator of Interferon Genes (STING); this interaction is required for STING’s ability to stimulate interferon expression, thus linking the Sec61 channel to the innate immune response (26). Sec61 with Sss1 can translocate proteins into the ER on its own, but the cytosolic domain of Sec61ß contacts secretory proteins prior to insertion in the cytosolic vestibule of the Sec61 channel and the presence of Sec61ß stimulates import (23, 27, 28). Recently, null mutations in human SEC61B were identified as a cause for polycystic liver disease (29). The mutations led to a complete biogenesis defect specific for polycystin-1, a large complex transmembrane protein with a weak secretory signal sequence (29). Collectively, these data have led to the view that Sbh1/Sec61ß mediates interaction of the Sec61 channel with other proteins in the ER membrane and that its cytosolic domain aids insertion of specific signal peptides or transmembrane domains into the Sec61 channel.

**FIGURE 1:**
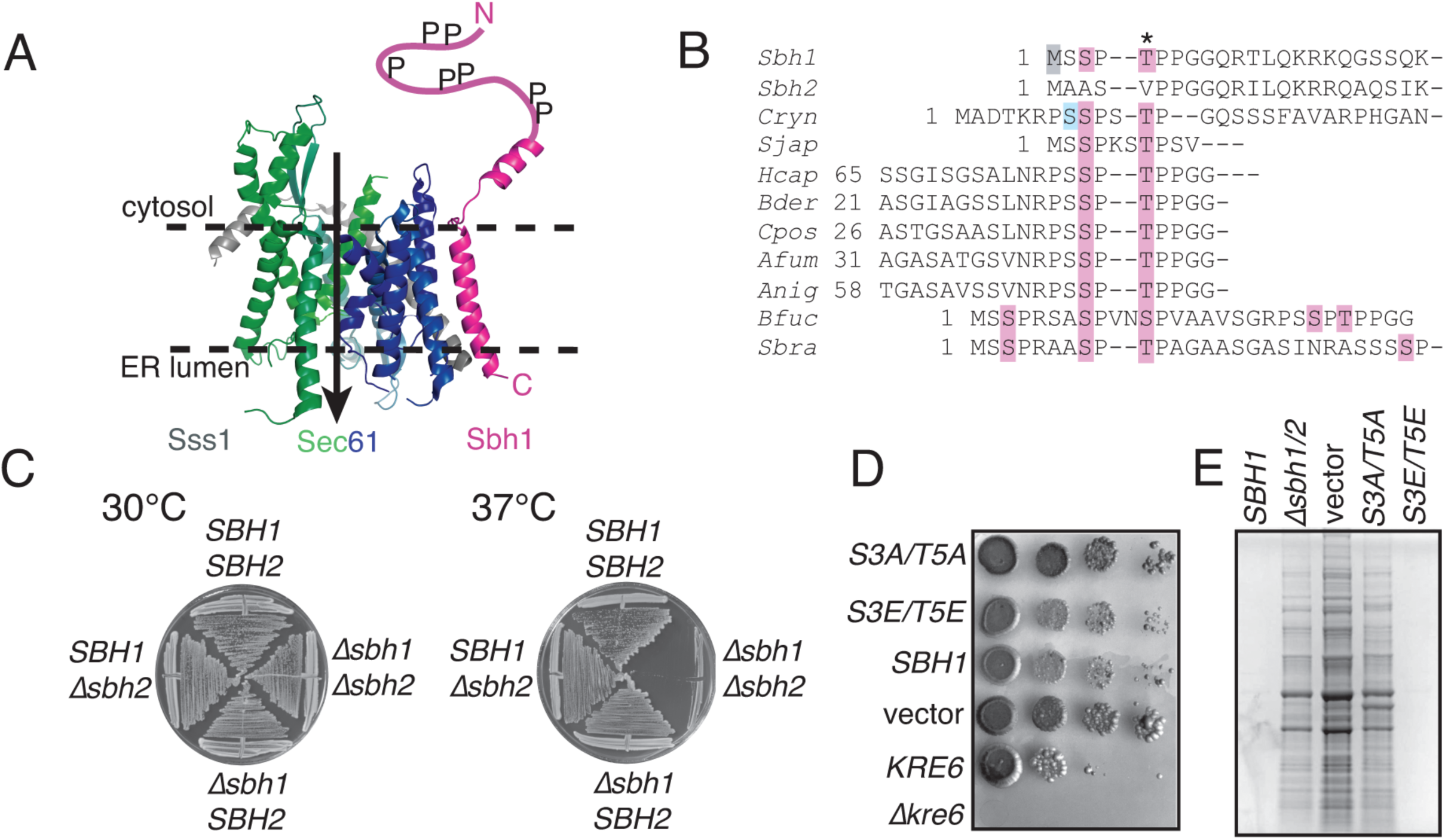
Sbh1 is required for cell wall integrity in *S. cerevisiae*. **A)** Structure of the Sec61 channel of *S. cerevisiae* (modified from 12; PDB 6ND1). The N-terminal half of Sec61 is shown in blue, C-terminal half in green, Sss1 in grey, and Sbh1 in magenta. The intrinsically unstructured N- terminal 38 amino acids of Sbh1 that were not visible in the cryo-EM structure were drawn in by hand. P, phosphorylation sites; dashed lines, position of the ER membrane; arrow, direction of protein transport. **B)** Alignment of the poorly conserved N-termini of Sbh1 proteins from *S. cerevisiae* (Sbh1, Sbh2) and the following pathogenic fungi: Cryn *Cryptococcus neoformans*, Sjap *Schizosaccharomyces japonicus*, Hcap *Histoplasma capsulatum*, Bder *Blastomyces dermatitidis*, Cpos *Coccidioides posadasii*, Afum *Aspergillus fumigatus*, Anig *Aspergillus niger*, Bfuc *Botryotinia fuckeliana*, Sbra *Sporothrix brasiliensis*. Except for Sbh2, only regions containing S or T residues with flanking prolines (pink) are shown. The conserved proline-flanked T5 in *S. cerevisiae* Sbh1 is indicated by an asterisk; other confirmed phosphorylation sites in *S. cerevisiae* Sbh1 are shown in grey; and the PKA site in *C. neoformans* Sbh1 is in blue. *Schizosaccharomyces cryophilus*, *Schizosaccharomyces octoporus*, *Candida albicans*, *Schizosaccharomyces pombe*, *Kluyveromyces lactis*, *Yarrowia lipolytica*, *Pichia pastoris*, and *Hansenula polymorpha* Sbh1 proteins were also screened, but do not contained potential proline-flanked phosphorylation sites. For full alignments of all Sbh1 protein sequences see Supplementary Figure 1. **C)** Wildtype *SBH1 SBH2 S. cerevisiae* and the indicated mutants were grown on YPD plates at the indicated temperatures for 3 days. **D)** The *Δsbh1Δsbh2 S. cerevisiae* strain was transformed with pRS415 without an insert (vector), wildtype *SBH1*, or the indicated phosphorylation site mutants and grown on SC without leucine containing 10 µg/ml calcofluor white at 30°C. A *Δkre6* mutant which is unable to grow on calcofluor white and its isogenic wildtype were included as controls. Note that these are in a different strain background compared to the *SBH1* strain. **E)** The indicated strains, including the untransformed *Δsbh1Δsbh2 S. cerevisiae* strain (*Δsbh1/2*), were grown to early exponential phase and cell-wall proteins were extracted by high pH/DTT, resolved by SDS-PAGE (size range shown is roughly 10-150 kD), and visualized by Coomassie staining.

Phosphorylation is a frequent biological mechanism for regulation of protein activity or protein- protein interactions and a systematic analysis identified 63 kinases that play a role in the pathogenicity of *C. neoformans* (30, 31). One of these, Protein Kinase A (PKA), regulates the secretion of a subset of virulence factors (32). Regulation of protein secretion may be important during physiological transitions such as cell differentiation or development. Indeed Sec61ß - the only subunit of the Sec61 channel that has been found to be phosphorylated so far - is important for polarization of epithelial cells, and for development of *Drosophila melanogaster* embryos (33, 34, 35, 36). Both mammalian Sec61ß and yeast Sbh1 are phosphorylated at multiple sites in various combinations (33, 21; phosphosite databases). The phosphorylation sites all lie in the intrinsically unstructured cytosolic domain, and are not positionally conserved, with one exception: the proline- flanked site at T5 in *S. cerevisiae* is present in mammals and some birds, although not in lower vertebrates, invertebrates, or other non-pathogenic yeast species. This suggests that this phosphorylation site evolved at least twice by convergent evolution and thus likely has an important function (Fig. 1A, asterisk Fig. 1B) (21).

The 63 kinases required for virulence of *C. neoformans* include many proline-directed kinases (31, 37). When we aligned the N-termini of Sbh1 proteins from non-pathogenic and pathogenic yeast, we noticed that - with the exception of *Candida albicans* - all pathogen Sbh1 proteins contain one or more potential proline-flanked phosphorylation site(s) in the N-terminal cytosolic domain, whereas Sbh1 proteins from non-pathogenic yeast species (*Schizosaccharomyces pombe, Kluyveromyces lactis, Yarrowia lipolytica, Pichia pastoris, Hansenula polymorpha*) and the *S. cerevisiae* homolog Sbh2 did not (Fig. 1B, pink; see Supplementary Figure 1 for full alignments). This led us to hypothesize that these phosphorylation sites in Sbh1 might be connected to virulence of the respective pathogenic species. In *C. neoformans* Sbh1 the proline-flanked phosphorylation sites are preceded by a potential PKA phosphorylation site (Fig. 1B, blue) and secretion of a subset of virulence factors in *C. neoformans* depends on PKA (32). We therefore decided to investigate the role of Sbh1 and its N-terminal phosphorylation sites in virulence of *C. neoformans*.

We found that, unlike in *S. cerevisiae,* mutations of the N-terminal Sbh1 phosphorylation sites (individually or in combination) do not affect *C. neoformans* growth in rich medium (YPD), and that even complete deletion of *SBH1* only inhibits growth when combined with high temperature and or cell wall stressors. We also observed that in infection-like conditions the deletion mutant fails to properly regulate protein secretion; we have identified the Sbh1-dependent substrates and identified common characteristics of their ER-targeting sequences. Notably, the affected proteins include multiple virulence factors and the mutant is consequently avirulent in a mouse model of infection. Together, our results show that Sbh1 is critical for secretion of a subset of virulence factors and other proteins during *C. neoformans* infection, and is therefore essential for its pathogenicity.

## RESULTS

### Sbh1 is required for cell wall integrity in *S. cerevisiae* and *C. neoformans*

We first looked to the best characterized system, *S. cerevisiae*. In this organism, both genes that encode the Sec61 channel beta subunit (*SBH1* and *SBH2*) are required for growth on YPD at 37°C (Fig. 1C) (17, 18, 21). *S. cerevisiae* lacking *SBH1* and *SBH2* are also resistant to calcofluor white (Fig. 1D), which suggests either reduced chitin in the cell wall or altered cell wall architecture.

Further, we observed increased extractability of proteins from the cell wall by alkaline buffer containing dithiotreitol, which also suggests structural defects in the wall (Fig. 1E). While mutation of the conserved Sbh1 phosphorylation site T5 to A on its own has no effect on *S. cerevisiae* growth or cell wall integrity (21), mutation of both proline-flanked phophorylation sites to A phenocopied the absence of Sbh1 in *S. cerevisiae*. Mutation of both residues to phosphomimetic E had no effect (Fig. 1D, E).

The phenotypes we observed in *C. neoformans* cells lacking their sole *SBH1* gene were quite distinct from those of *S. cerevisiae*. Specifically, they showed only subtle temperature sensitivity at 37°C on YPD (Fig. 2A, left panel) and no sensitivity to osmotic or cell wall stressors at 30°C (Fig. 2A). At 37°C, however, the mutant cells were hypersensitive to SDS and barely grew on calcofluor white (Fig. 2A, lower panels; Fig. 2B, top two rows); they were also more sensitive than wild-type cells to cell wall digestion by lysing enzymes (Fig. 2C) and showed increased sensitivity to oxidative stress (1 mM H_2_O_2_; Fig. 2D, top two rows), which is encountered in the setting of infection. Overall, we conclude that Sbh1 is required for cell wall integrity in both the ascomycete *S. cerevisiae* and the basidiomycete *C. neoformans*, although specific manifestations of this requirement differ (see Discussion).

**FIGURE 2:**
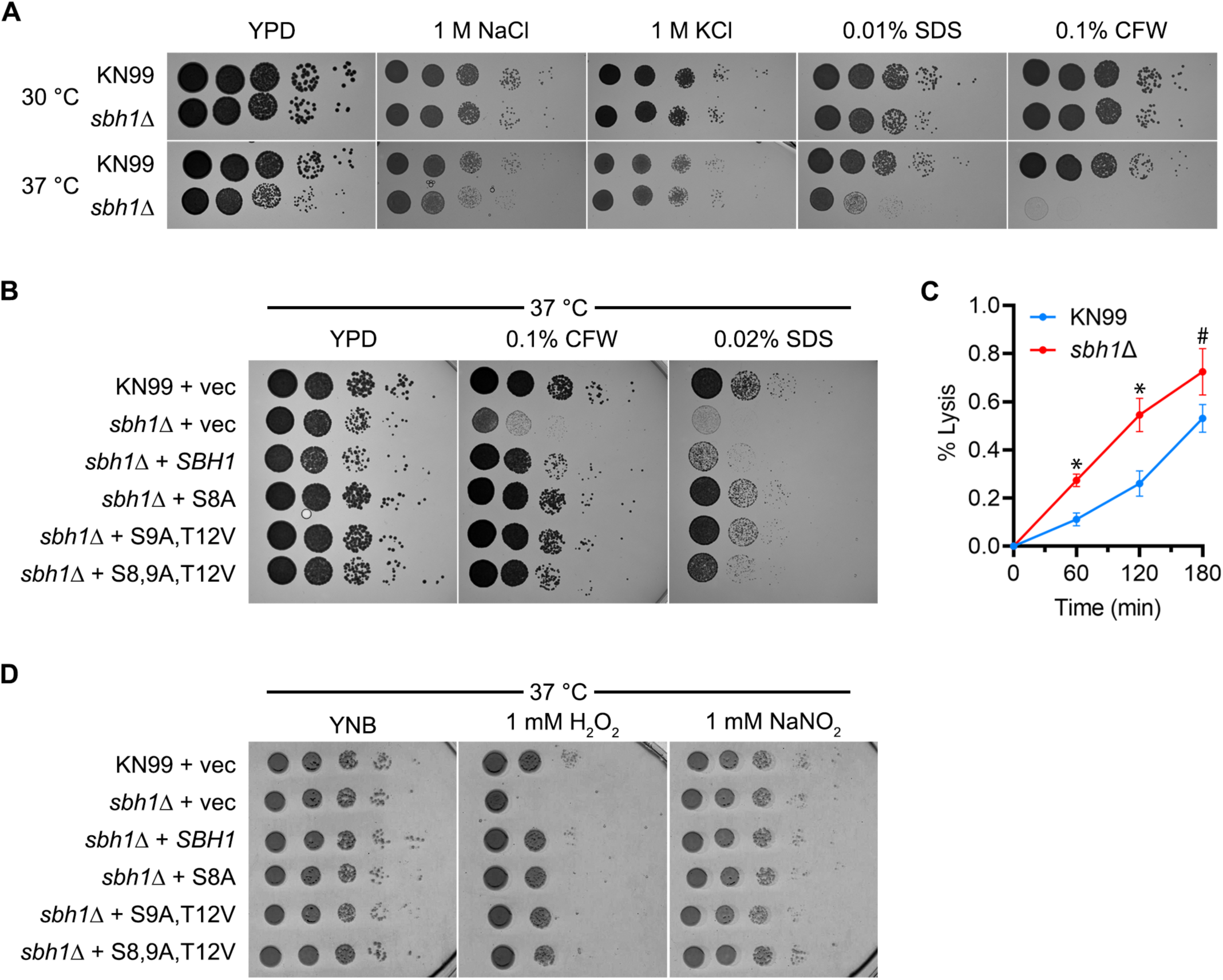
Sbh1 is required for cell wall integrity at high temperature in *C.neoformans*. **A)** Serial 10-fold dilutions of wild-type (KN99) and *Δsbh1* cells were grown at the temperatures shown on YPD alone or containing 1 mg/ml calcofluor white (CFW) or 0.02% SDS (SDS). **B)** Wild type (KN99) or *sbh1Δ* mutant cells were transformed with vector alone (vec) or plasmids encoding HA-tagged Sbh1 (SBH1), or Sbh1 with mutations in the indicated phosphorylation sites and then diluted as in panel A and grown at the indicated conditions. **C)** Cells of the strains indicated were treated with lysing enzymes (see Methods) for the times shown and then assessed for % lysis with two-way ANOVA with mixed-effects analysis. * indicates P<0.05, # indicates P=0.25. **D)** The indicated strains were grown as above in the presence of 1 mM H_2_O_2_ or 1mM NaNO_2_ on YNB (pH 5) at 37°C. All experiments are representative of at least three biological replicate studies.

### Sbh1 N-terminal phosphorylation sites do not affect cell wall integrity in *C. neoformans*

As discussed above, we hypothesized that the N-terminal phosphorylation sites in cryptococcal Sbh1 were important for protein function. To test this, we first engineered a plasmid to express wildtype *SBH1* modified with a C-terminal HA tag. The tagged protein effectively complemented the sensitivity of *sbh1Δ* mutants to cell wall stressors (CFW and SDS, Figure 2B). We then modified this expression plasmid to encode Sbh1 that was altered to eliminate either the putative PKA site (Ser8 to Ala), two proline-flanked phosphorylation sites (Ser9 to Ala and Thr12 to Val), or all three sites. All of these polypeptides were expressed at similar levels, except for the version with a sole PKA site mutation, which was slightly less abundant (Supplementary Figure 2). We found that each of the mutated proteins fully complemented growth of *sbh1Δ* cells in the presence of CFW and also restored growth at 37 °C in the presence of SDS (Fig. 2B). Interestingly, cells expressing mutant versions of Sbh1 grew slightly better on YPD and were more resistant to SDS than those expressing the WT protein (Fig. 2B). Each mutant protein also completely complemented the defective growth of the *sbh1Δ* mutant in the presence of oxidative stress (Fig. 2D). Together, these results suggest that, surprisingly, these phosphorylation sites are not required for Sbh1 function, as assessed by its role in cell wall integrity.

### Sbh1-dependent secretory and transmembrane proteins are induced in conditions that mimic the host environment

Sbh1 contacts signal peptides in the cytosolic vestibule of the Sec61 channel, so it can both guide secretory proteins into the channel and control their access to the secretory pathway. To address whether biogenesis of specific secretory or transmembrane proteins was compromised in the absence of *C. neoformans* Sbh1, we analyzed the proteomes of wild-type and *sbh1Δ* cells grown in either rich medium (YPD) at 30°C or in DMEM at 37°C and 5% CO_2_. The latter, termed 37D5 below, was chosen to mimic key features of the complex host environment. We detected no significant differences between wildtype KN99 and *sbh1Δ* strains in YPD at 30°C (Fig. 3), suggesting no critical role for Sbh1 in those conditions. In 37D5, however, we discovered a set of proteins that was specifically induced in wild-type cells, but not in the *sbh1Δ* mutant (Fig. 3, Supplementary Table 1). This set included 32 proteins without secretory pathway targeting sequences, 21 transmembrane (TM) proteins and 68 proteins with signal peptides (SP) or uncleaved signal anchors (SA) (Supplementary Table 1). When we analyzed their ER-targeting sequences we found that the transmembrane domains were slightly more hydrophobic than average for transmembrane proteins in *C. neoformans*; many also had a strong positively charged patch in close proximity to one side of the TM domain (Fig. 4A; Supplementary Table 2). Some of the transmembrane domains of the Sbh1-dependently induced proteins also contained a high number of prolines and/or glycines in addition to the polybasic stretch adjacent to the transmembrane domain (Supplementary Table 2). The Sbh1-dependent signal peptides were more heterogeneous, but most displayed reduced C-region polarity (Fig. 4B; Supplementary Table 2). In addition, they were suboptimal in various ways which therefore defied bioinformatic analysis: some had limited hydrophobicity, some contained high numbers of proline and glycine which interfere with alpha- helix formation, and many had a charge bias problem (either no charge on either side of the hydrophobic core, or a positive charge C-terminal instead of N-terminal of the hydrophobic core, or multiple positive charges at the N-terminus of the mature part of the protein) (Supplementary Table 2). Because signal peptides insert as alpha-helices into the hydrophobic lateral gate of the Sec61 channel with the positively charged N-terminus towards the cytosol, any of these features will reduce translocation efficiency into the ER. We suggest that Sbh1 is required for the biogenesis of specific secretory and transmembrane proteins under host-like conditions. Our model is that because the targeting sequences of these proteins are suboptimal for Sec61 channel insertion, they rely on the enhanced insertion efficiency provided by the presence of Sbh1 (Fig. 4C) (23).

**FIGURE 3:**
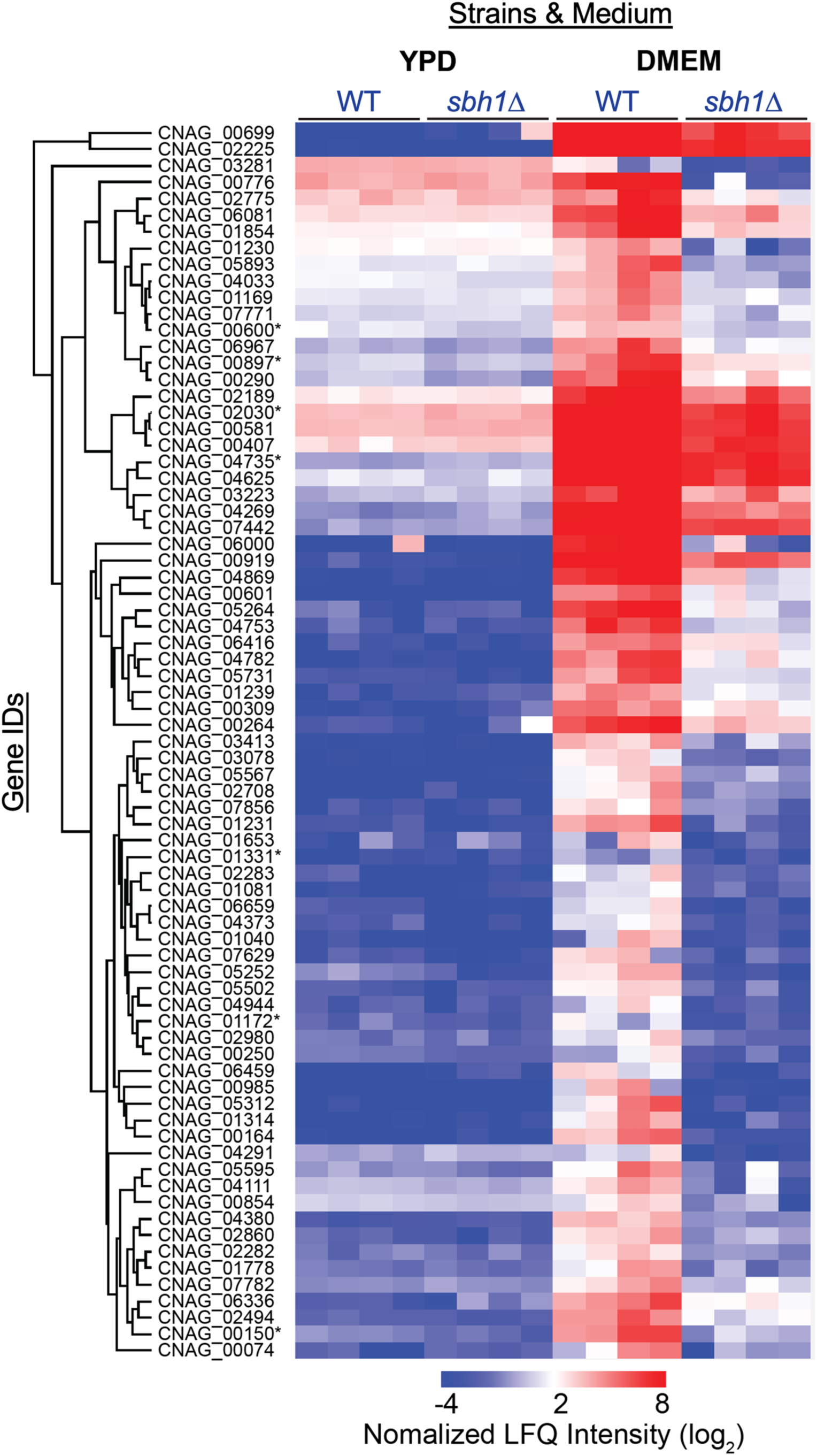
*C. neoformans* Sbh1 is required for the induction of secretory and transmembrane proteins in conditions that mimic the host. Proteomic analysis of wildtype and *sbh1Δ C. neoformans* grown in YPD at 30°C or in 37D5. Samples were analyzed in quadruplicate. Gene IDs of known virulence factors are indicated by asterisks.

**FIGURE 4:**
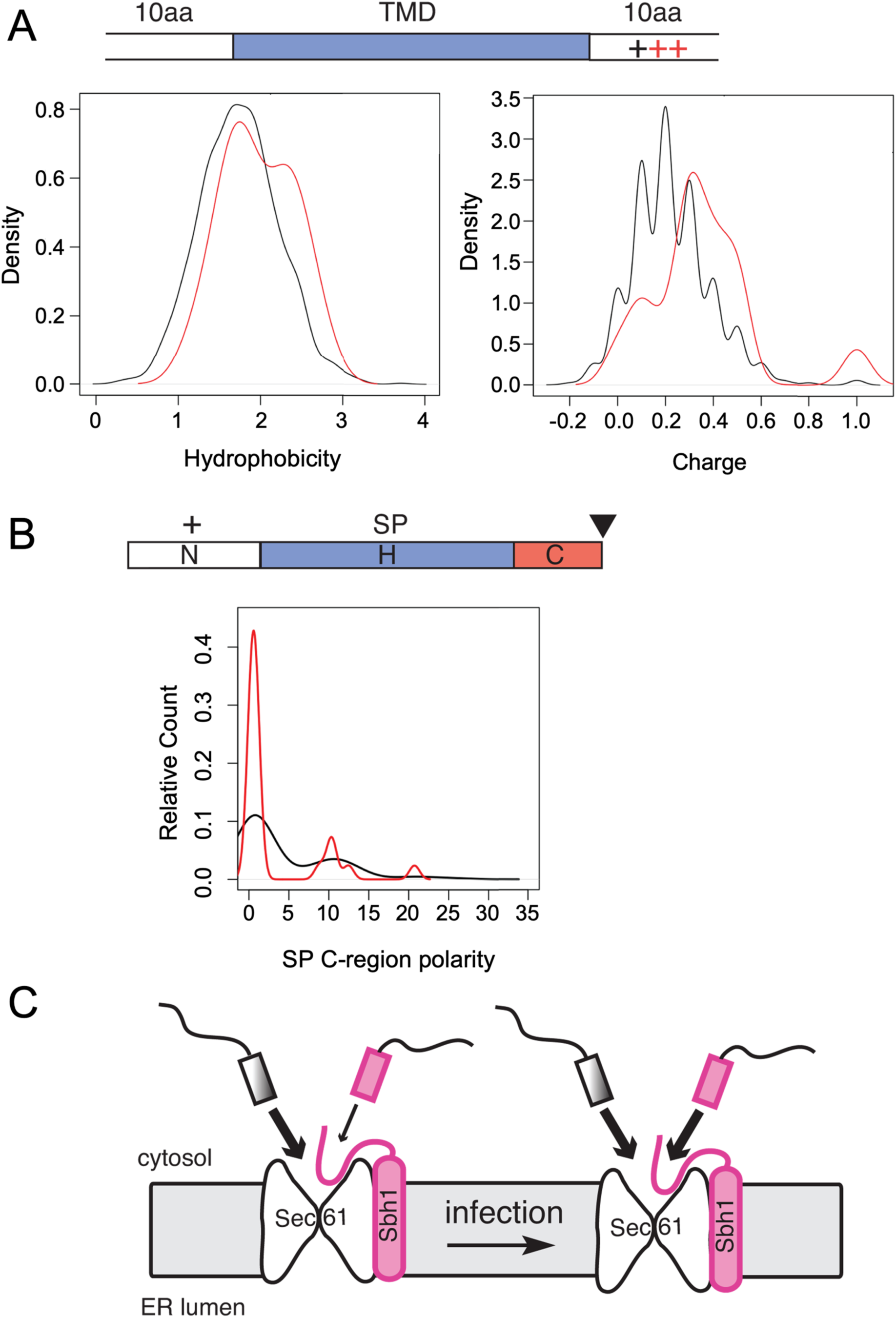
Secretory and transmembrane proteins that require Sbh1 for induction contain specific ER-targeting sequences. **A)** Characterization of Sbh1-dependent transmembrane domains. Top, schematic of a typical transmembrane domain (TMD) with features specific for Sbh1-dependent TMDs highlighted in red. Hydrophobicity (left) and charges within 10 amino acids of TMDs (right) were analyzed for TMDs of Sbh1-dependently induced proteins (candidates, red) compared to all transmembrane proteins in the *C. neoformans* proteome (black). **B**) Characterization of Sbh1-dependent signal peptides. The schematic on top shows a typical signal peptide (SP) with features specific for Sbh1-dependent SPs highlighted in red. Black triangle: signal peptidase cleavage site. SPs of Sbh1-dependently induced secretory proteins (red) were analyzed for N-region charge, H-region hydrophobicity and length, and C-region polarity and compared to all signal peptides in the *C. neoformans* proteome (black). The only difference found was in C-region polarity, shown in the graph. **C)** Model for Sbh1 function in *C. neoformans*. Under rich medium conditions, proteins with Sbh1-dependent ER targeting sequences (magenta) play only a minor role for vegetative growth (left), and most proteins imported into the ER have Sbh1- independent targeting sequences (shaded grey). During infection (right), however, Sbh1-dependent ER import (magenta) becomes essential.

### Sbh1 is essential for virulence of *C. neoformans*

The proteome analysis presented above shows that Sbh1 regulates the secretion of proteins that are induced by host-like conditions. This process is likely important for pathogenesis, as supported by our observation that *sbh1Δ* cells have a reduced ability to survive within host macrophages (Figure 5, panels A and B). To directly test the role of Sbh1 in virulence, we used a well-established mouse model for cryptococcal infection. At 14 days after intranasal infection we observed a striking reduction in both lung and brain fungal burden in *sbh1Δ*-infected animals compared to wild type (Figure 5, panels C and D). Consistent with this finding, mice infected with wild-type cryptococci succumbed to infection within 18 days, while mutant-infected animals remained healthy and grew normally for at least 10 weeks (Figure 5, panels E and F). We conclude that *SBH1* is essential for *C. neoformans* virulence.

**FIGURE 5:**
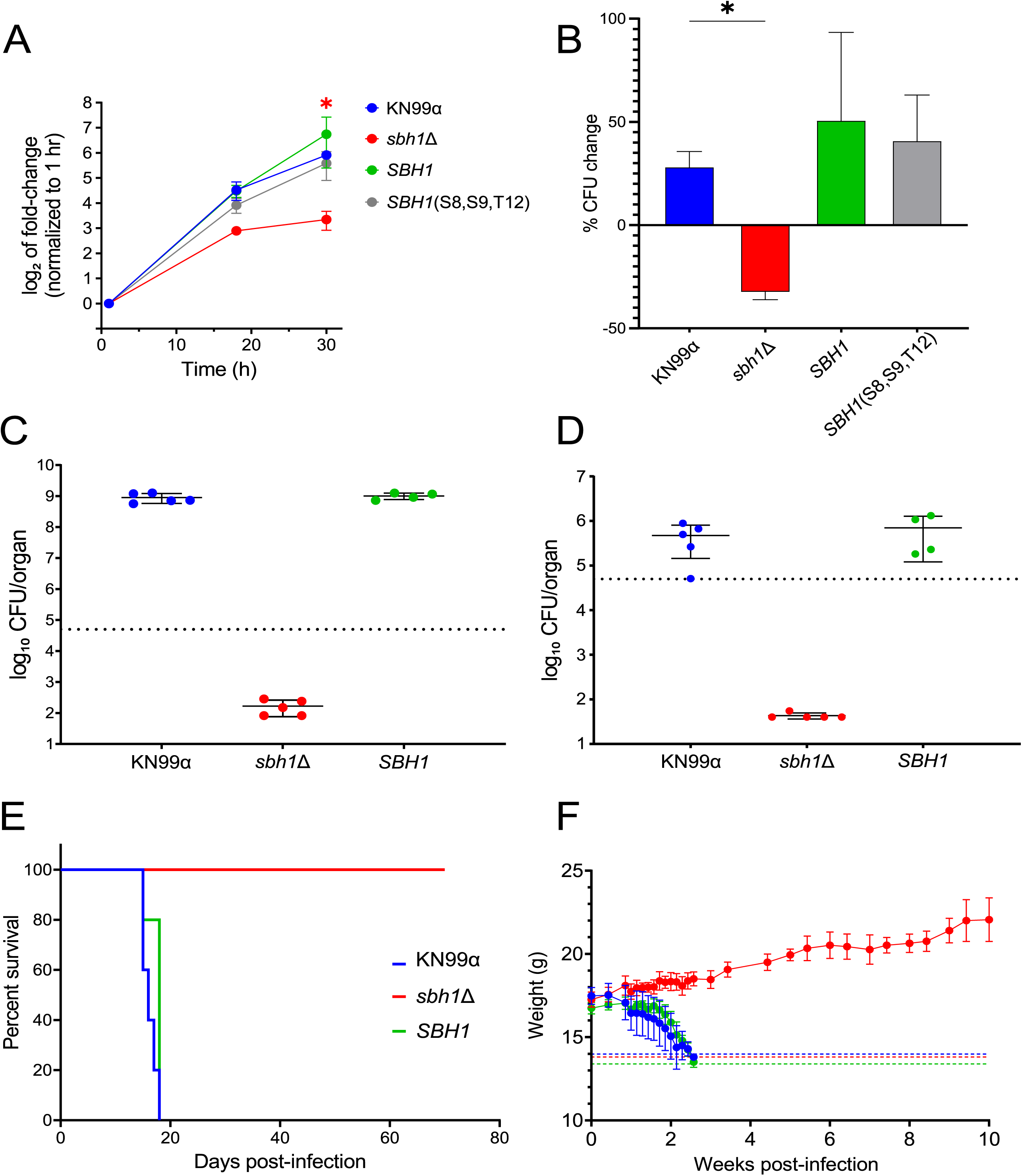
The *C. neoformans sbh1*Δ mutant is defective in growth in conditions that mimic the host or within host cells and is avirulent in mouse infection. (A) Fungi of the indicated strains were incubated in tissue culture medium (5% CO_2_, 37 °C) for 1, 18, or 30 h and then plated on YPD for enumeration of colony-forming units (CFU; shown normalized to the 1 h CFU value). Shown are mean and SEM values from 4 independent experiments. (B) Cells were incubated for 1 h with differentiated THP-1 cells (MOI = 1), during which time fungi were engulfed by the host cells. Samples were then vigorously washed to remove free cryptococci and host cells were lysed to measure CFU either immediately or after 30 h of incubation. % change was calculated as [(Final CFU – Initial CFU) / (Initial CFU)] x 100]. Shown are mean and SEM values from 2 independent experiments. For (A) and (B), * indicates p < 0.05 for *sbh1Δ* compared to KN99. (C, D) Total fungal burden at 14 days post-infection with the indicated strains, measured in lung (C) and brain (D). Each symbol represents colony forming units (CFU) for one mouse; mean and standard deviation are also shown. Dotted line, original inoculum. (E) Mouse survival after intranasal infection with 5x10^4^ cells of KN99 wildtype (solid line), *sbh1Δ* (dashed line) or *sbh1Δ* complemented with wildtype *SBH1* (dotted line). (F) Weight (mean and SD) of the mice from Panel E over time after infection. Any mouse that decreased to less than 80% of its initial weight (indicated by dashed lines) was humanely sacrificed and considered not to survive the infection.

## DISCUSSION

In this work we sought to identify factors regulating secretion of virulence-relevant proteins in the human pathogen *C. neoformans*. We specifically investigated the role of the non-essential ER protein translocation channel subunit Sbh1. Since the Sbh1 cytosolic domain contacts secretory signal sequences in the cytosolic vestibule of the channel prior to insertion into the channel it can potentially aid or restrict protein import into the secretory pathway of specific proteins under specific growth conditions. This may be regulated by phosphorylation as Sbh1 in all species contains multiple phosphorylation sites which are only partially positionally conserved (Fig. 1A, B).

In *S. cerevisiae*, absence of *SBH1* and its paralog *SBH2* causes temperature sensitivity, although the ER translocation defects at the restrictive temperature are modest for most substrates investigated (Fig. 1C; 38, 18). We hypothesized that this phenotype could relate to defects in the cell wall for several reasons: other translocation mutants exhibit cell wall defects (39); wall synthesis relies on secretory proteins; and alterations in this process often yield a temperature-sensitive phenotype (40). We investigated the effect of *sbh1* mutants on cell wall integrity by assessing growth on calcofluor white (CFW; Fig 1D) and extractability of cell wall proteins by high pH/dithiotreitol treatment (Fig. 1E). CFW binds to chitin in the yeast cell wall and is toxic (41). *KRE6* encodes a glucosyl hydrolase required for beta-1,6-glucan synthesis; the deletion mutant is hypersensitive to CFW and is shown as a control. Cells lacking Sbh1 were slightly resistant to CFW (Fig. 1D), indicating a reduced amount of chitin in the cell wall, and displayed increased extraction of cell wall proteins (Fig. 1E).

*S. cerevisiae* cell wall integrity also specifically required the two proline-flanked N-terminal phosphorylation sites at S3 and T5; mutation of these residues to non-phosphorylatable A phenocopied the cell wall defects of the deletion mutant (Fig. 1D, E). The *sbh1 S3E/T5E* mutant behaved like wildtype (Fig. 1D, E). Proline-flanked phosphorylation sites in Sbh1 of other yeast species occur primarily in human pathogens (Fig. 1B), which suggested to us that phosphorylation at these sites may play a regulatory role in virulence. The absence of *SBH1* in *C. neoformans* resulted only in a very modest temperature sensitivity at 37°C (Fig. 1C vs. 2A), although at high temperature *sbh1Δ C. neoformans* also displayed a cell wall defect (Fig. 2A-C). In contrast to *S. cerevisiae*, however, mutation of the N-terminal proline-flanked phosphorylation sites of *C. neoformans* Sbh1 individually or in combination did not phenocopy the cell wall defect (Fig. 2B). As protein kinase A has been shown to be critical for *C. neoformans* virulence, and *C. neoformans* Sbh1 also contains a PKA site (Fig. 1B, blue), we also mutated this site alone or in combination with the proline-flanked sites, but again saw no effect on cell wall integrity (Fig. 2B) (32).

Surprisingly, the altered phosphorylation sites slightly enhanced cell growth, especially notable on SDS (Fig. 2B). Taken together, our data suggest that Sbh1 is important for efficient secretion of one or more enzymes or precursors for the cell wall in both *S. cerevisiae* and in *C. neoformans*, but that the role of the N-terminal proline-flanked phosphorylation sites in this process differs in the two organisms. Alternatively, the differential effect of the phosphorylation site mutations may reflect their distinct cell wall compositions: whereas the *S. cerevisiae* cell wall is formed by ß-1,3-glucan, ß-1,6-glucan, chitin, and mannoproteins, in the *C. neoformans* cell wall most chitin is deacylated to chitosan, and *α*-1,3-glucan is a major cell wall component (42, 10).

We next asked which proteins were affected by absence of *SBH1* in *C. neoformans*. Using a proteomic approach we identified a set of secretory and transmembrane proteins that were induced under 37D5 conditions in the wild-type strain, but not in the *sbh1Δ* mutant (Fig. 3; Supplementary Tables S1 and S2). Many of the membrane proteins in this set were characterized by a strong polybasic patch adjacent to the first transmembrane domain (e.g. CNAG_1854, CNAG_5502, CNAG_6416), while the signal peptides of the secretory proteins were more heterogeneous, and had less polar C-regions than average (Supplementary Table S2; Fig. 4A, B). This is reminiscent of the ER targeting signals identified by Ziska and colleagues in mammalian cells whose ER import was dependent on the Sec61 channel-interacting protein Sec63 and the ER-lumenal Hsp70 chaperone BiP (43). In *S. cerevisiae* the Sec63 complex composed of Sec63, Sec62, Sec71, and Sec72 promotes posttranslational protein import into the ER through the Sec61 channel by stabilizing the lateral gate in a partially open conformation (44, 12). It also plays a role in membrane protein topogenesis in both yeast and mammalian cells (45, 46). Sec63 is a polytopic membrane protein with an ER-lumenal J-domain that activates ATP-hydrolysis of the ER-lumenal chaperone BiP (Kar2 in yeast) (47). Kar2 contributes to protein import into the ER either by directly binding to the incoming protein or by promoting its folding (48, 49). As proteins dependent on Sbh1 in *C. neoformans* and proteins dependent on Sec63/BiP in mammalian cells have similar characteristics, our data suggest that Sbh1 and the Sec63 complex have overlapping functions in promoting ER import of specific proteins.

**Table 1.** Sbh1-dependent proteins involved in cryptococcal virulence. These proteins meet the following criteria: their expression in 37D5 was at least two-fold higher in WT compared to sbh1Δ cells; they contain an SP or TM domain; and they are required for normal C. neoformans virulence in mouse infections. Note that this is likely a subset of such proteins, as most cryptococcal proteins have not been tested for roles in virulence; other candidates that meet the first two criteria are listed in Supplementary Table 1. Proteomics results are shown as the ratio of expression in WT versus sbh1Δ cells in 37D5. RNA-seq results are shown as the ratio of gene expression at 24 h versus 0 h in 37D5 (61). Expression of a subset of these genes (CNAG_04735, CNAG_02030, CNAG_00897) was measured by qPCR; these studies confirmed that it is not affected by loss of SBH1 (Supplementary Figure 3).

| Gene ID | Gene name | Protein function | SP or TM | WT/*sbh1Δ* proteomics [log2-fold] | Virulence citations | 24 h/0 h RNA-seq |
| --- | --- | --- | --- | --- | --- | --- |
| CNAG_00150 |  | serine endoptidase, cell wall | SP1-23 | 5.24 | 53 | 1.68 |
| CNAG_01653 | *CIG1* | cytokine-inducing glycoprotein | SP1-19 | 4.00 | 68 | 385.34 |
| CNAG_01172 | *PBX1* | beta-helix containing protein | SP1-18 | 3.68 | 55,53,56 | 0.78 |
| CNAG_04735 | *MPR1* | extracellular metalloproteinase (Mep1) | SP1-19 | 3.44 | 53, 59 | 122.79 |
| CNAG_00897 | *SKN1* | glucosidase | TM134-58 | 3.04 | 57 | 3.43 |
| CNAG_02030 |  | glyoxal oxidase | SP1-21 | 2.79 | 53 | 0.97 |
| CNAG_00600 | *CAP60* | capsule-associated protein | SP1-33 | 2.65 | 58 | 1.97 |
| CNAG_01331 |  |  | SP1-18 | 2.62 | 53 | 13.45 |

Signal peptides generally insert into the Sec61 channel with their N-terminus towards the cytosol and their N-terminal positive charge (Fig 4B, ’+’) contributes to this orientation (50). Many of the Sbh1-dependent signal peptides we identified in *C. neoformans* have no charge bias towards the N- terminus; in some, the charge bias is even reversed (Supp. Table S2). Sbh1 may help orient such signal peptides in the cytosolic vestibule of the Sec61 channel with the help of its cytosolic domain. The polar C-region of signal peptides contains the signal peptidase cleavage site (50). Most of the Sbh1-dependent signal peptides we identified have a very short C-region (Supp. Table S2) which may limit signal peptidase access unless the signal peptide is positioned accurately by Sbh1. In order to efficiently insert into the Sec61 channel the signal peptides of secretory proteins require an alpha-helical hydrophobic region (Fig 4B, H region, blue) (50). Many of the Sbh1-dependent signal peptides of *C. neoformans* contain multiple helix-breaking P or G residues within their hydrophobic helical core regions, and some have very short or very long hydrophobic H-regions (Supp. Table 2). These features would make it difficult for signal peptides to stably insert into the Sec61 channel lateral gate. Overexpression of an N-terminally truncated Sbh1 lacking most of its cytosolic domain complements the temperature-sensitivity and at least some of the translocation defect in *Δsbh1 Δsbh2 S. cerevisae* (18). Together with our data, this suggests that the Sbh1 transmembrane domain stabilizes a conformation of the Sec61 lateral gate that favours insertion of the helical hydrophobic core of signal peptides, even if they are suboptimal.

That Sbh1-dependent secretion is critical for virulence is illustrated by both the reduced intracellular fitness of *sbh1Δ C. neoformans* in macrophages and the effects in mice. In contrast to infection with KN99 wildtype *C. neoformans*, which resulted in a high burden in the lungs and death of all mice by day 18 post infection, all mice infected with *sbh1Δ C. neoformans* survived for 70 days and even gained weight (Figure 5), showing that the mutant strain was avirulent.

We hypothesize that the striking defect in virulence of the *sbh1* mutant reflects reduced secretion of multiple groups of proteins. For a few examples, these could include proteins required for fungal growth in the host environment (e.g. metabolic genes or proteins involved in cell wall synthesis), needed for virulence factor production, or directly involved in host damage. These categories may also interact; for example, mutations that alter cell wall glycan synthesis also result in aberrant formation of the capsule (7, 8, 51, 52). To explore this idea, we reviewed the polypeptides whose secretion is most impaired in the absence of Sbh1 (Supplementary Table 1), focusing on proteins with signal or transmembrane sequences that had been previously implicated in cryptococcal virulence. We identified eight such proteins, which had exhibited infectivity defects in a large-scale screen (53) and/or been individually studied and shown to contribute to virulence (Table 1). One of them, Cig1, has also been detected in the blood of mice infected with *C. neoformans*, leading to the suggestion that it could serve as a biomarker of infection (32). Notably, all of the corresponding genes are expressed in human CSF during cryptococcal meningitis, most at higher levels than in rich medium (Table 1 and (54)). Two proteins in this group (Pbx1(55, 56) and Skn1 (57)) act in cell wall synthesis, so their reduced secretion could contribute to the wall defects and sensitivity to host-induced stress that we observed in *sbh1* mutant cells. Another is required for capsule display (Cap60 (58)), although its biochemical function has not been characterized, and two others are proteases (CNAG_00150 and Mpr1 (59)). Many fungal proteases damage host tissues during infection (60) and Mpr1 further acts in cryptococcal crossing of the blood-brain barrier (59), which is required for cryptococcal dissemination to the brain and subsequent lethality. Supporting a role in infection for this group of proteins, six of the corresponding genes are expressed more highly in 37D5 than in rich medium (one other is modestly decreased); this pattern is particularly striking for *CIG1* and *MPR1*, whose expression levels increase over 100-fold (Table 1) (61). Notably, the evidence that these proteins are important for cryptococcal virulence derives from studies of individual gene deletion strains, which completely lack one of these target proteins. The situation in the *sbh1* mutant is different, in that many proteins are secreted less efficiently, but none (besides Sbh1) are absent. Nonetheless, the impaired secretion of multiple proteins with key roles in virulence, together with others in the functional groups above, provides abundant reason for the inability of *sbh1* to cause disease in a mammalian host.

We conclude that *C. neoformans* Sbh1 is important for biogenesis of virulence factors whose specific ER-targeting sequences make these proteins dependent on Sbh1 for ER import (Fig. 4C). Recently, Sec61beta has also been shown to be critical for infection by the important fungal crop pathogen *Magnaporthe oryzae*, suggesting a more general role for Sbh1/Sec61beta in fungal infection (62). Since the intrinsically unstructured N-terminal part of the Sbh1 cytosolic domain is not conserved between yeast and mammals, our work may suggest a new specific target for developing drugs against this important human pathogen.

## MATERIALS AND METHODS

### S. cerevisiae methods

*S. cerevisiae* strains are detailed in the Supplemental Methods and were grown on either yeast peptone dextrose (YPD) or synthetic complete medium without leucine, with or without 10 µg/ml calcofluor white (Sigma), at 30°C or 37°C, for 3 days. Cell wall proteins were extracted from cells in early exponential phase in 100 mM Tris-HCl, pH 9.4, 10 mM DTT at 37 °C, and TCA-precipitated for SDS-PAGE analysis as detailed in the Supplemental Methods.

### *C. neoformans* strains, growth conditions, and plasmids

All strains were in the *C*. *neoformans* serotype A strain KN99*α* background, as detailed in the Supplemental Methods, and were grown at 30°C on YPD with 100 µg/mL of nourseothricin or 100 µg/mL G418 as appropriate. After plasmids expressing the native *SBH1* gene or version with 5 HA epitopes at the C-terminus were shown to complement phenotypic defects of the *sbh1* mutant, the tagged version was mutagenized to modify N-terminal motifs and electroporated into KN99*α* and the *sbh1*Δ strains as described in (63) and detailed in the Supplemental Methods.

Because of the heterogeneity in copy number associated with plasmids in *C. neoformans*, at least three independent colonies were picked and tested for each construct; all exhibited the same behaviors.

### Phenotypic assays and mass spectrometry

Serial dilution plate assays and sensitivity to hypotonic lysis after exposure to lysing enzyme were assessed as detailed in the Supplementary Methods and (64). Total cellular proteome analysis was performed as in (65, 66) and the Supplementary Methods. Results were compared to the reference *C.neoformans* reference proteome and deposited in the PRIDE partner repository for the ProteomeXchange Consortium with the data set identifier: PXD013894. A Student’s *t*-test was performed to identify proteins with a significant differential expression (*p-value* < 0.05) (S_0_ = 1) between samples employing a 5% permutation-based FDR filter.

### Infection assays

Fungal survival after engulfment by THP-1 (ATCC #TIB-202) cells *in vitro* was assessed as in (67), with minor modifications detailed in the Supplemental Methods. For *in vivo* studies, 4-6 week-old female C57BL/6 mice were intranasally inoculated with 5x10^4^ cryptococcal cells and either followed by weight for survival studies or sacrificed at set times for organ burden measurement, as detailed in the Supplemental Methods. All animal studies were reviewed and approved by the Animal Studies Committee of Washington University School of Medicine and conducted according to the National Institutes of Health guidelines for housing and care of laboratory animals.

## ACKNOWLEDGMENTS

We thank William Allen, Bristol University, for help with Fig. 1A; Christina Soromani, Andrea Ludes, and Nicole Spitzlberger (all former students of the Römisch lab) for data shown in Fig. 1C, D; Alyssa R. Brunsmann, Daphne Ko, and Liza Loza for qPCR studies; and Jussi Jäntti, VTT Helsinki, for *S. cerevisiae* strains and plasmids.

KR was funded by core funding of the Saarland University. Work in the Doering lab was supported by NIH grants AI140979 and AI135012 to TLD. FHS also received support from NIH T32 grant AI007172 and a Postdoctoral Enrichment Program Award from the Burroughs Wellcome Fund. JGM is supported by the Canadian Foundation for Innovation (JELF #38798) and the Canadian Institutes for Health Research (Project Grant). VH was supported by Deutsche Forschungsgemeinschaft grant He3875/15-1. The funders had no role in study design, data collection and interpretation, or the decision to submit the work for publication.

## DATA AVAILABILITY

Proteomics .Raw data files are available in the PRIDE (Proteomics Identification Database): Project ID: PXD013894

**Password:** 8ZiMfbKG

## SUPPLEMENTARY INFORMATION

### SUPPLEMENTAL METHODS (word doc)

**TABLE S1.**
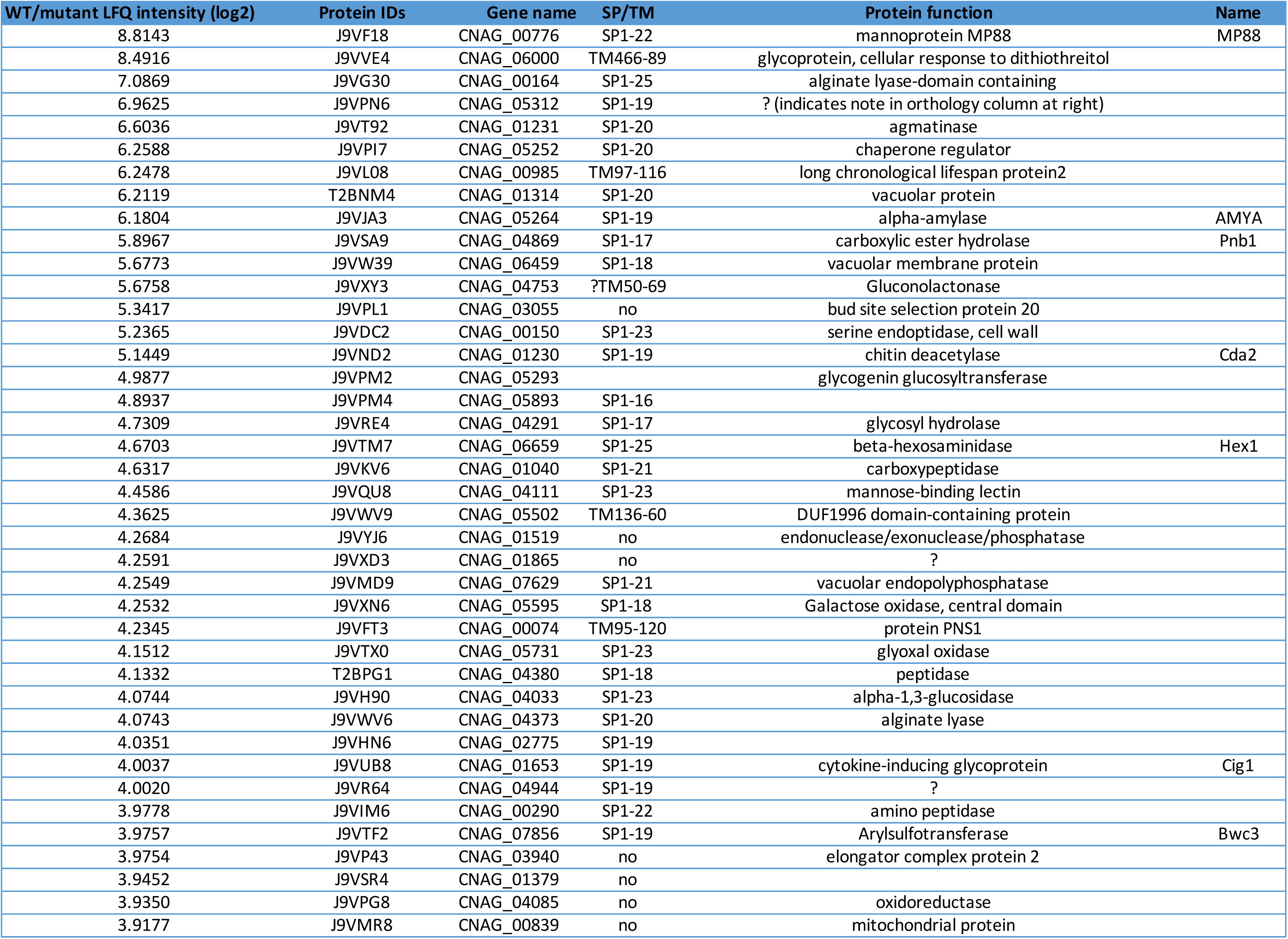
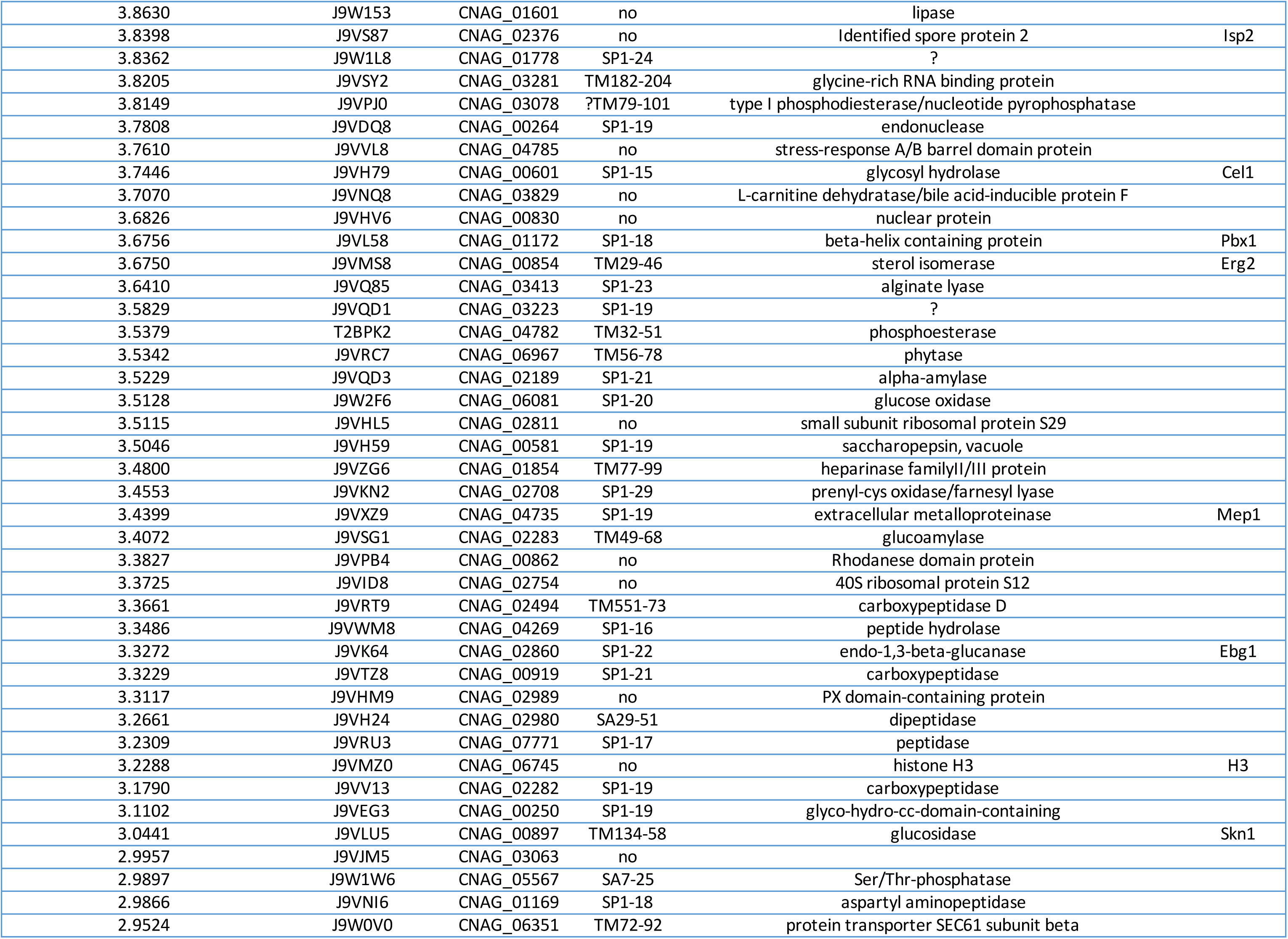
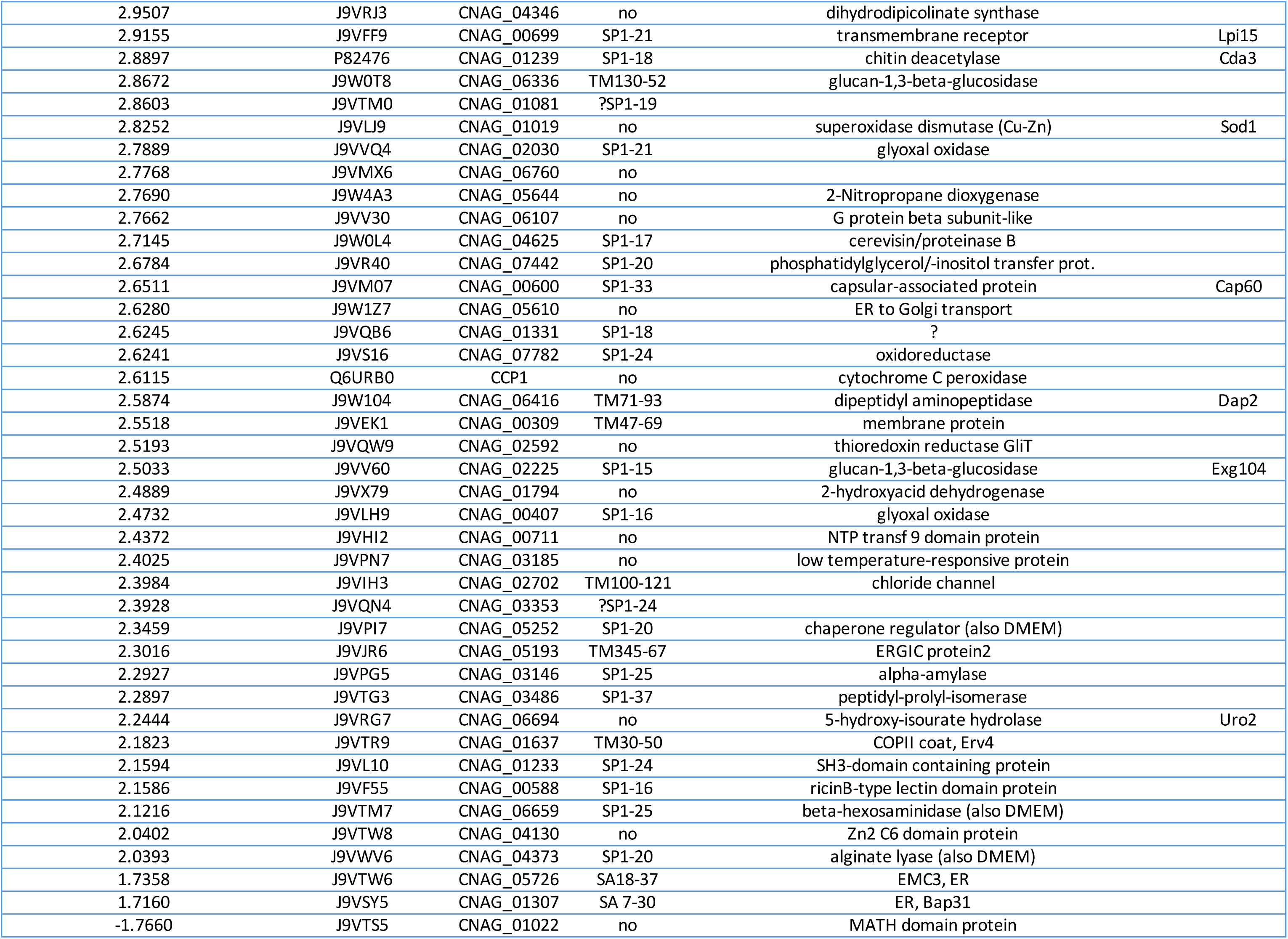
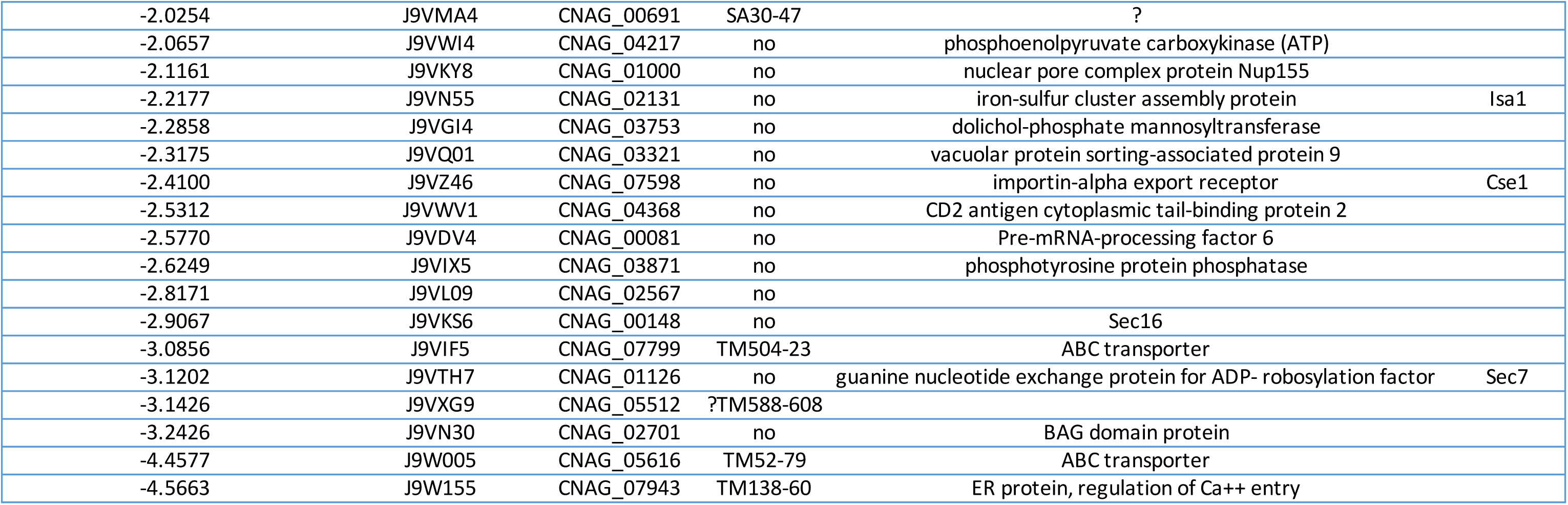
(Excel file): Protein expression analysis in KN99 wild-type and sbh1Δ C. neoformans grown in YPD at 30°C or DMEM at 37°C and 5% CO_2_. Proteins are ordered by degree of Sbh1-dependence (log_2_ of wild-type/sbh1Δ LFQ intensity). Where present, ER targeting sequences are indicated by amino acid numbers. SP, signal peptides; SA, uncleaved signal anchors; TM, first transmembrane domain. Ambiguous predictions due to limited hydrophobicity (TM) or suboptimal signal peptide parameters (SP) are indicated by ’?’.

**TABLE S2.**
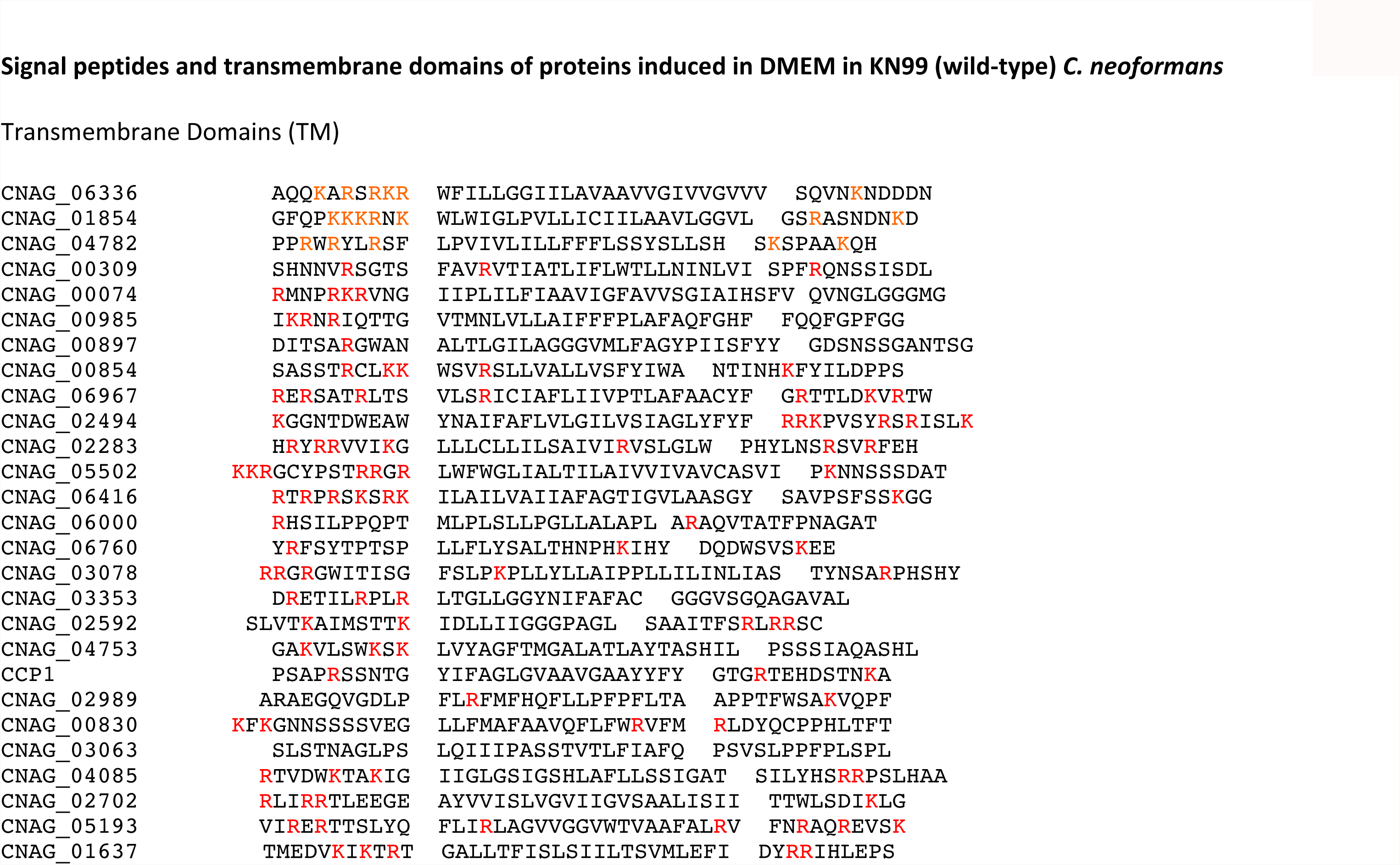
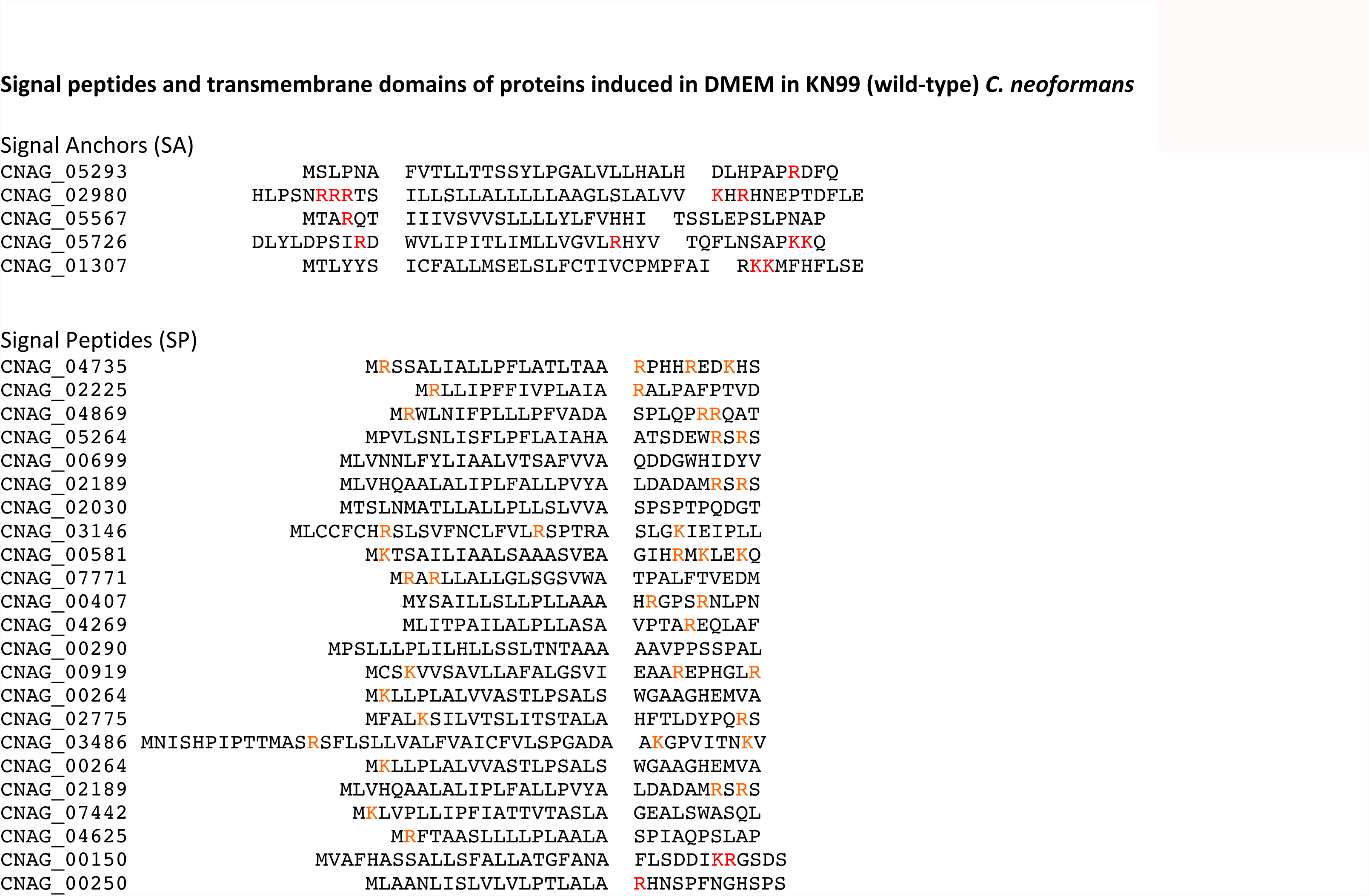
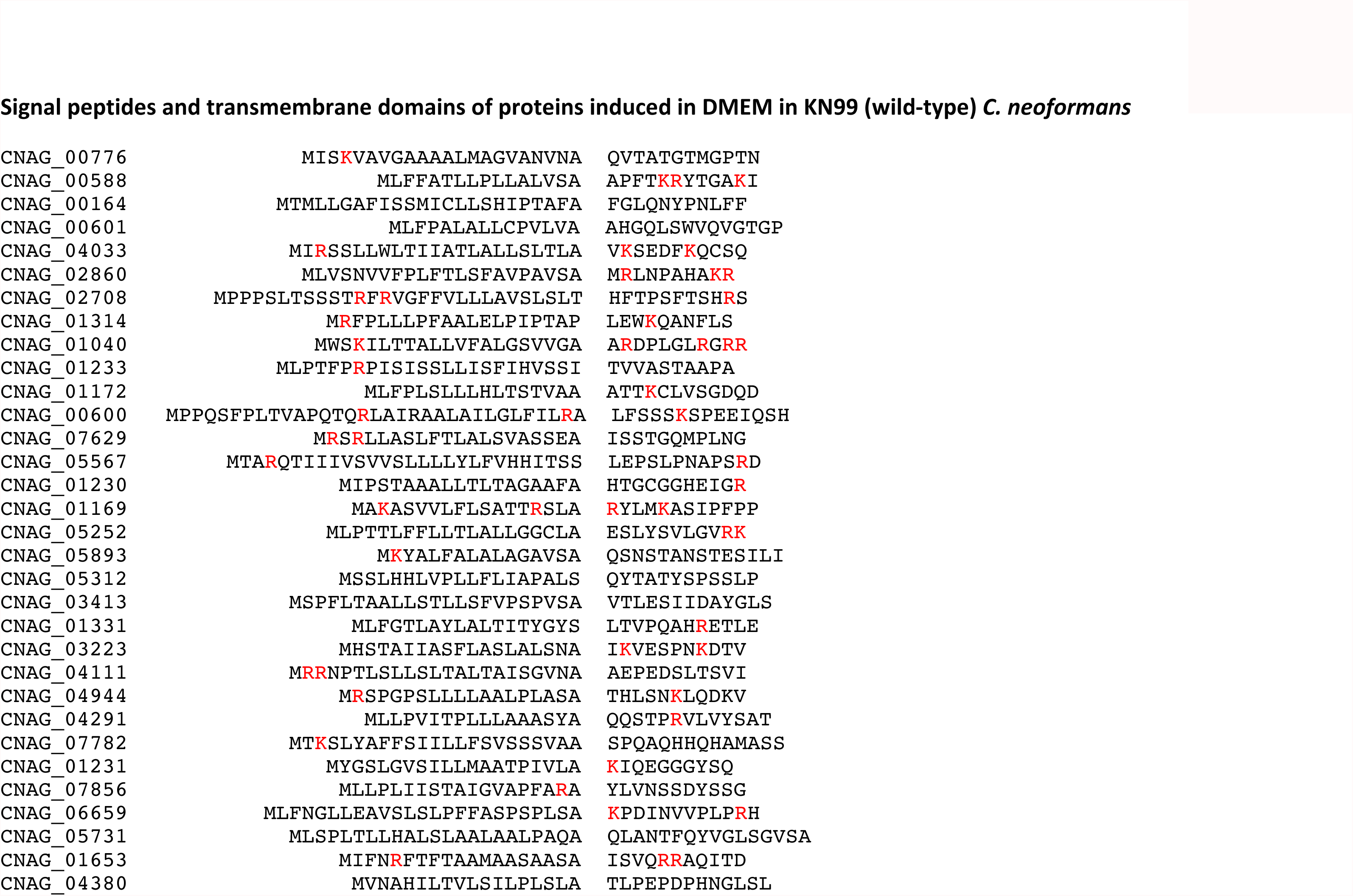
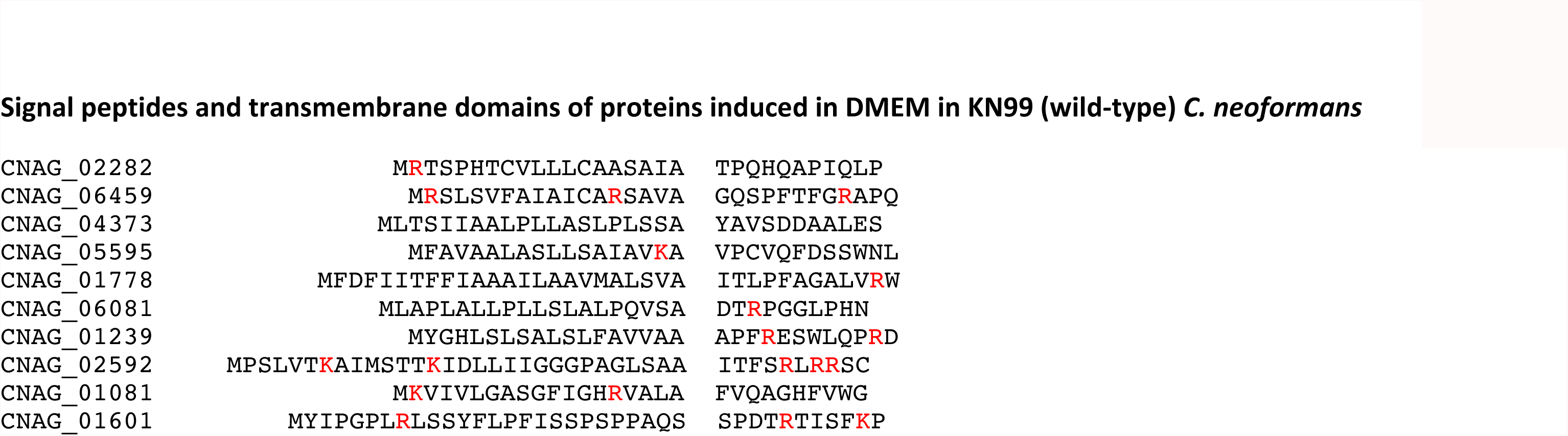
(word doc): Signal peptides and transmembrane domains of proteins induced in DMEM in KN99 wildtype *C. neoformans*. Amino acid sequences of TM domains, signal peptides, and signal anchors are shown including the flanking 10 amino acids on either side for TM or the first 10 amino acids of the mature domains for SP. Positively charged amino acid residues are highlighted in red.

**SUPPLEMENTARY FIGURE 1:**
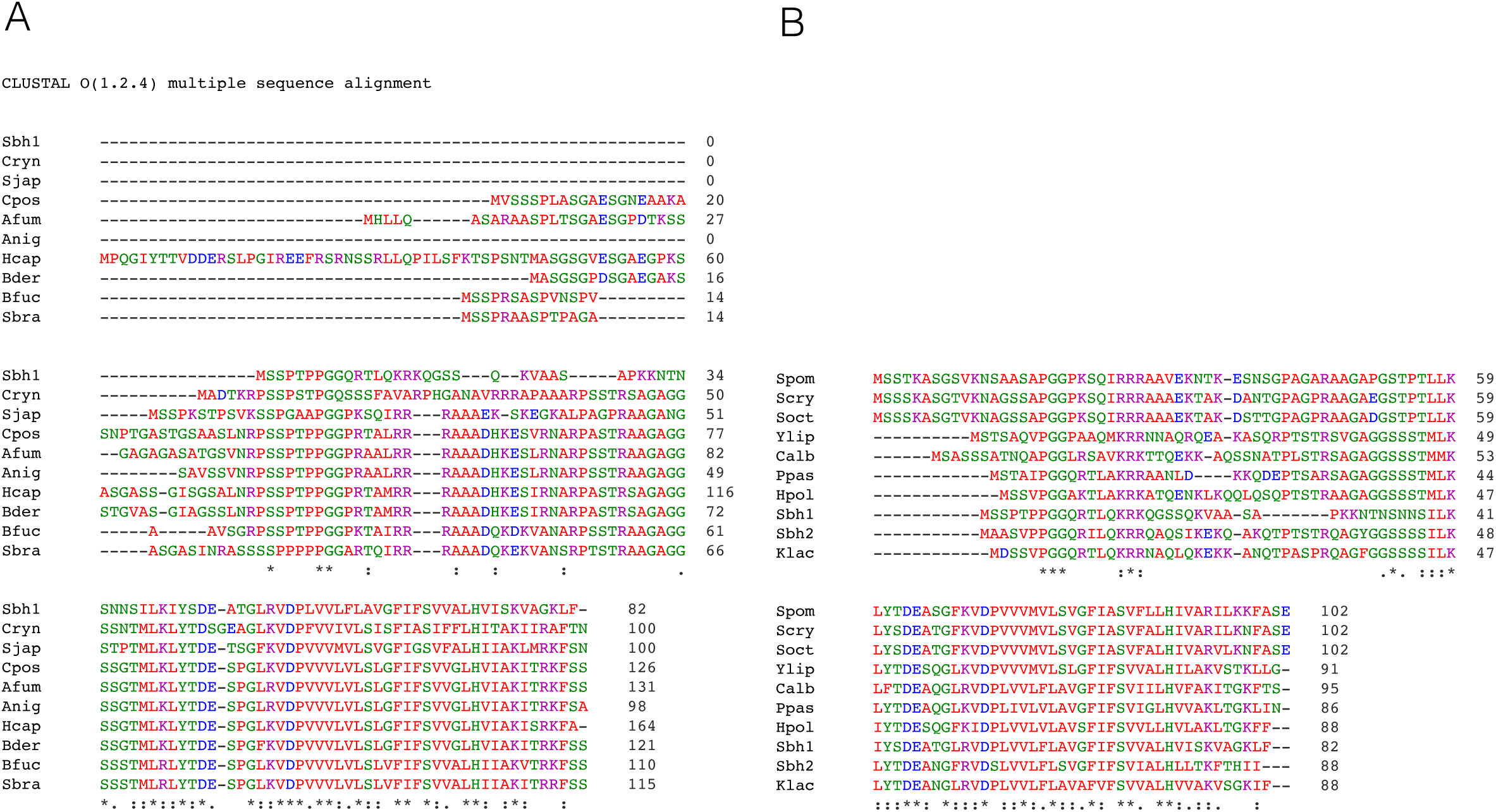
Alignments of full Sbh1 amino acid sequences. Protein sequences were extracted from Uniprot and aligned using Clustal Omega. (A) Sbh1 orthologues with N-terminal, proline-flanked S or T. Sbh1 = *S. cerevisiae*, proline-flanked N-terminal phosphorylation sites highlighted in yellow; Cryn *Cryptococcus neoformans*, Sjap *Schizosaccharomyces japonicus*, Hcap *Histoplasma capsulatum*, Bder *Blastomyces dermatitidis*, Cpos *Coccidioides posadasii*, Afum *Aspergillus fumigatus*, Anig *Aspergillus niger*, Bfuc *Botryotinia fuckeliana*, Sbra *Sporothrix brasiliensis*. (B) Sbh1 orthologues without N-terminal, proline-flanked S or T. *S. cerevisiae* Sbh1 is shown for comparison; Sbh2 = *S. cerevisiae* paralogue without N-terminal, proline-flanked S or T; Scry, *Schizosaccharomyces cryophilus*, Soct *Schizosaccharomyces octoporus*, Cal *Candida albicans*, Spom *Schizosaccharomyces pombe*, Klac *Kluyveromyces lactis*, Ylip *Yarrowia lipolytica*, Ppas *Pichia pastoris*, Hpol *Hansenula polymorpha*.

**SUPPLEMENTARY FIGURE 2:**
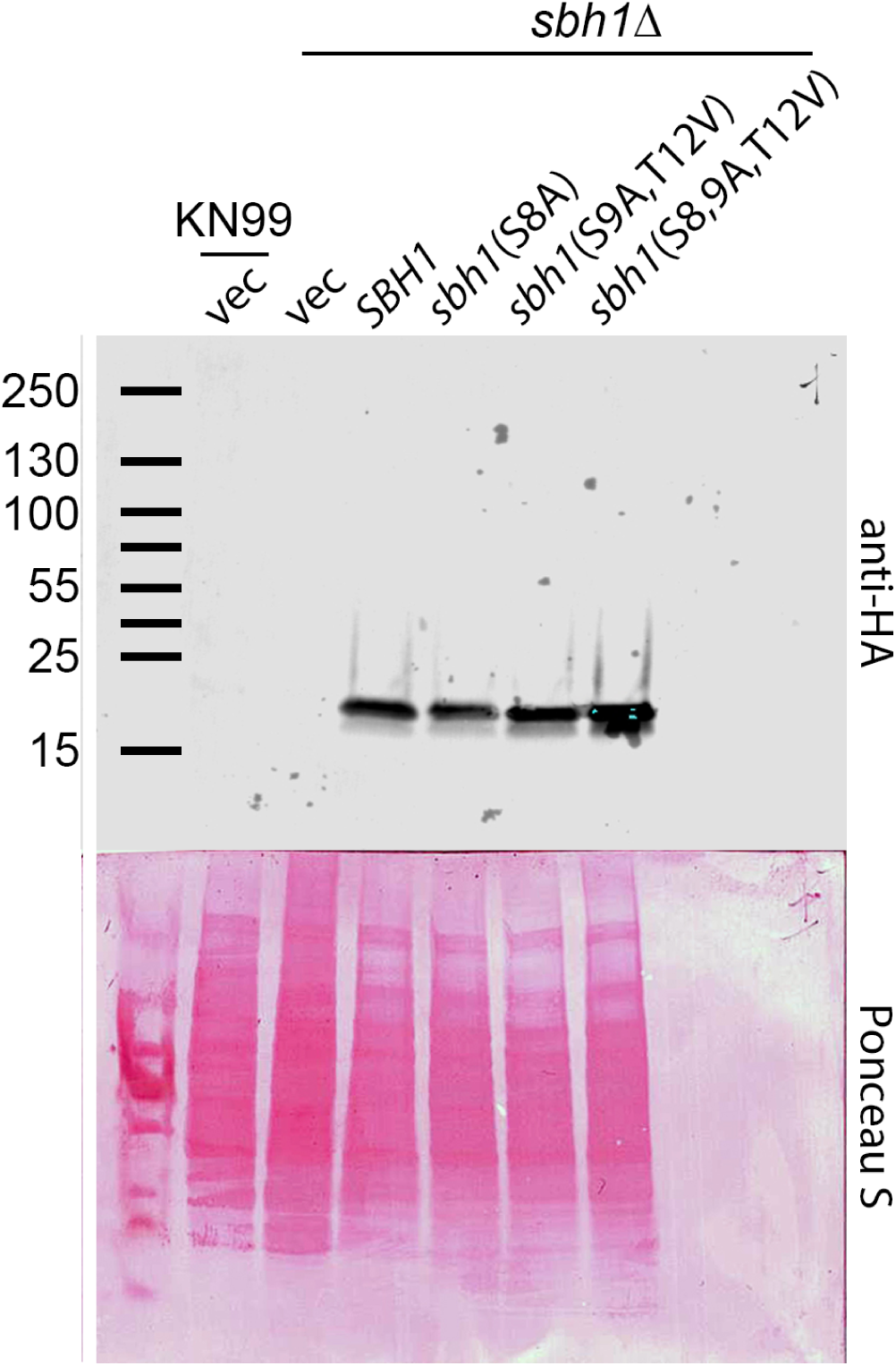
Expression of wild type and phosphorylation mutants of Sbh1. Shown is an anti-HA immunoblot of wildtype or *sbh1*Δ cryptococcal cells expressing either vector alone (vec) or the indicated HA-tagged proteins. 20 ug of lysate were resolved on a 4 – 20% gradient gel and transferred onto PDVF membrane, which was stained with Ponceau S. This blot is representative of three independent experiments. The numbers and lines on the left represent the protein ladder. The Sbh1-5XHA protein is ∼16kDa.

**SUPPLEMENTARY FIGURE 3:**
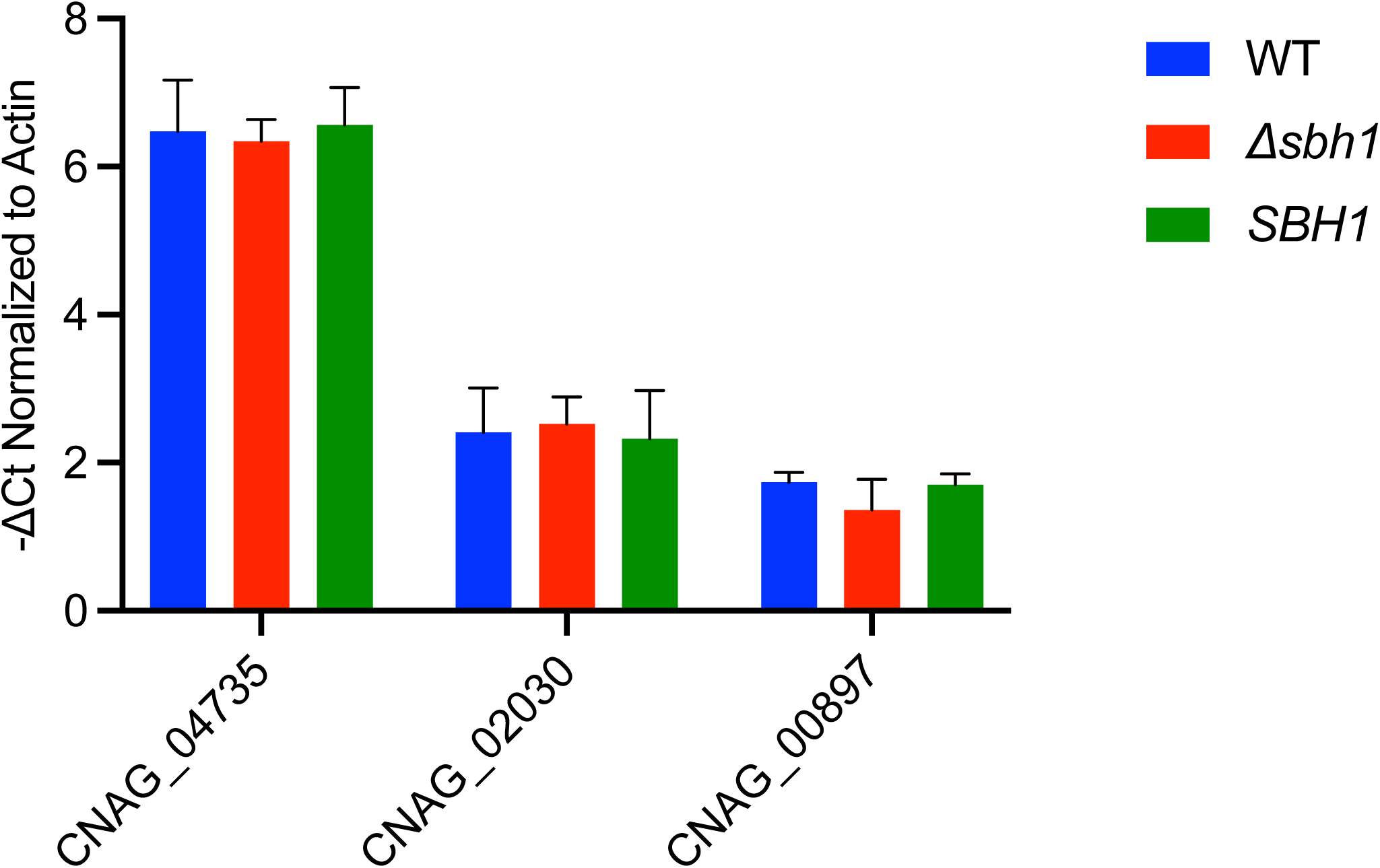
qRT-PCR analysis of selected genes in WT, *sbh1* mutant, and complemented strains grown in conditions that mimic the host. (DMEM, 37°C, 5% CO_2_, 24 h). For each gene indicated, the plot shows mean and standard deviation of results from three biological replicates, each done in technical triplicate and normalized to *ACT1* expression. There were no significant differences in gene expression for any pairwise comparison between strains (all P > 0.2 by unpaired T-test).

## SUPPLEMENTAL METHODS

### *S. cerevisiae* strains and growth conditions

NY179 (SBH1 SBH2 MAT a leu2-3,112 ura3-52), H3223 (MAT a KanMx::sbh1 leu2-3,112 ura3-52 GAL^+^), H3203 (MAT a HphMx::sbh2 leu2-3,112 ura3-52 GAL^+^), and H3231 (MAT a KanMx::sbh1 HphMx::sbh2 leu2-3,112 ura3-52 GAL^+^) were gifts from Jussi Jäntti and used to characterize Sbh1p phosphorylation sites (Toikkanen et al., 1996; Feng et al., 2007). SBH1 cloned into pCR2.1-TOPO plasmid was kindly provided by Jussi Jäntti (Helsinki University, Finland). The QuickChange Site-Directed Mutagenesis Kit (Stratagene, UK) was used to introduce single or multiple base mutations in SBH1. Substitutions were verified by sequencing. Mutated sbh1 was excised from pCR2.1-TOPO plasmids using EcoRV/BamHI, and subcloned into pRS415. For growth assays cells were grown on YPD or synthetic complete medium without leucine and with or without 10 µg/ml calcofluor white (Sigma) at 30°C or 37°C for 3 days.

### *S. cerevisiae* cell wall extraction

Cells were grown in synthetic complete medium without leucine to early exponential phase and 10 OD_600_ harvested by centrifugation. Cells were resuspended in 100 µl 100 mM Tris-HCl, pH 9.4, 10 mM DTT, and incubated for 15 min at 37°C shaking at 800 rpm in a Thermomixer (Eppendorf). Samples were chilled for 1 min on ice, followed by sedimentation of the cells in a refrigerated Eppendorf centrifuge at maximal speed for 1 min. 90 µl of each supernatant were transferred to a new tube, carefully avoiding the pellet, and centrifuged again as above. 80 µl of each supernatant were transferred to a fresh tube and proteins precipitated by adding an equal volume of ice-cold 20 % trichloroacetic acid. Samples were mixed thoroughly and incubated on ice for 30 min. Proteins were sedimented by centrifugation at 4°C for 15 min and the pellets washed with ice-cold acetone, dried, and resuspended in 20 µl 2x SDS sample buffer by heating for 10 min to 95°C with agitation. Proteins were resolved on 4-14% Bis-Tris gels (NuPage) run in MOPS buffer and revealed by Coomassie staining.

### *C. neoformans* strains and growth conditions

All strains used were in the *C*. *neoformans* serotype A strain KN99a background. The *sbh1*Δ deletion in this background used here was obtained from a library of partial gene deletions (generated by the Madhani group (Liu et al, 2008)) and available from the Fungal Genetics Stock Center at Kansas State University, Manhattan, KS) and confirmed by PCR. Fungal stocks were maintained at -80°C and grown at 30°C on yeast peptone dextrose (YPD) media with antibiotics as appropriate (100 µg/mL of nourseothricin (clonNAT, WERNER BioAgents, Germany) or 100 µg/mL G418 (Geneticin, Life Technologies, USA)).

### *SBH1* plasmid construction

For plasmid construction, the *SBH1* gene with its promoter and terminator sequences was amplified and cloned into ApaI/SpeI-digested pIBB103 (Skowyra and Doering, 2012). In parallel, a similar fragment lacking the terminator sequences and stop codon was amplified and cloned into ApaI/SpeI-digested pFS5XHA (pIBB103 containing 5 HA epitopes followed by the *TRP1* terminator; this study). After both versions were confirmed to complement phenotypic defects, the tagged version was used for all subsequent analyses. This construct was used as template for mutagenesis of the N-terminal motifs of *SBH1* by using overlapping primers containing the codon changes and the forward and reverse primers used above. The resulting product was digested, cloned into pFS5XHA, and the whole insert sequenced to confirm mutations. These plasmids, together with an empty plasmid control, were electroporated into KN99a and the *sbh1*Δ strains as detailed below.

### Electroporation of *C. neoformans*

Plasmid transformation was done by electroporation as described in (Skowyra and Doering, 2012). In brief, *Cryptococcus* cells patched on YPD agar plates with appropriate antibiotics were transferred into 5 ml of YPD liquid medium with antibiotic and cells were grown overnight at 30°C with shaking at 250 rpm. The overnight culture was diluted to an OD_600_ of 0.05 in 50 ml of fresh YPD medium (note that this OD_600_ value was measured in a multiplate reader and was not corrected for pathlength). After two doubling times the cells were transferred into conical tubes and sedimented, washed twice with cold dH_2_O, and resuspended thoroughly in 50 ml cold EB buffer (10 mM Tris-HCl, pH 7.5, 1 mM MgCl_2_, and 270 mM Sucrose) with 4 mM DTT. After incubation on ice for 15 min, the cells were pelleted, washed with 50 ml EB buffer without DTT, and resuspended in 1 ml EB buffer for counting. Cells were adjusted to 300 million cells per 100 µl, mixed with at least 1 µg of DNA (in no more than 1/10 of the total volume) and the mixture transferred into ice-cold 2 mm gap cuvettes. A Biorad electroporator was used with the following settings: V = 500v, R = infinity ohms, C = 25µF; the time constant should be 15 – 25 ms. Immediately after electroporation, 900 µl YPD without antibiotics was added and the cell suspension was transferred to a culture tube and incubated at 30°C for 3-4 h for recovery before various dilutions were spread on YPD plates with antibiotics to select transformants. Because of the heterogeneity in copy number associated with plasmids in *C. neoformans*, at least three independent colonies were picked and tested for each construct. All exhibited the same behaviors.

### Phenotypic analysis

For stress plating, strains to be tested were grown overnight in YPD with antibiotic as appropriate, diluted to an OD_600_ of 0.05 in fresh YPD, and grown for two doublings. The cultures were then adjusted to 2 x 10^9^ cells/ml, serially diluted (10-fold) and 5-µl aliquots were spotted onto the phenotyping plates described in the text. These plates did not include antibiotics because in combination with stressors they inhibit cell growth. Plates were incubated at 30°C and 37°C for 3-4 days.

The assay for sensitivity to lysing enzymes was adapted from (Gerik et al., 2005). Briefly, overnight cultures of wild type and mutant strains in YPD medium were washed once with PBS and once with citrate buffer (10 mM, pH 6.0). Cells were resuspended in citrate buffer and the OD_600_ was normalized to 1.0/ml with the same buffer. 1 ml aliquots were then subjected to centrifugation and the pellets (one per time point) were resuspended in a solution of lysing enzyme (final concentration 25 mg/ml; Sigma, L-1412) and incubated at 37°C with gentle agitation. At each desired timepoint one aliquot per strain was sedimented and resuspended in the same volume of distilled water (in the absence of lysis the OD_600_ should remain ∼1.0). The assay was performed independently three times.

### Mass spectrometry analysis

For total cellular proteome analysis, cells were collected by centrifugation at 3,500 rpm for 10 min and the pellets were washed twice with cold PBS before being subjected to acetone precipitation and enzymatic digestion (Geddes et al., 2016; Ball and Geddes-McAlister, 2019). For secretome analysis, supernatant fractions from *C. neoformans* WT and *sbh1*Δ cell cultures were collected and filtered using 0.22 µM syringe filters and subjected to in-solution trypsin digestion (Ball and Geddes-McAlister, 2019). Digestion was stopped by the addition of 10% v/v trifluoroacetic acid (TFA) and the acidified peptides were desalted and purified according to the standard protocol (Rappsilber et al., 2007). Approximately 50 µg of sample was loaded onto StageTips and stored at 4°C until LC-MS/MS measurement.

For LC-MS/MS measurement, samples were analyzed by nanoflow liquid chromatography on an EASY-nLC 1200 system (ThermoFisher Scientific) on-line coupled to a Q Exactive HF-X quadrupole orbitrap mass spectrometer (ThermoFisher Scientific). An in-line 75 µm x 50 cm PepMap RSLC EASY-Spray column filled with 2 µm C18 beads (ThermoFisher Scientific) separated peptides using linear gradients from 3% to 20% Buffer B (80% acetonitrile) over 18 min and from 20% to 35% Buffer B over 31 minutes, followed by a steep 2 min ramp to 100% Buffer B for 9 min in 0.1% Formic acid at a constant flow of 250 nl/min.

Raw files were analyzed together using MaxQuant software (version 1.6.0.26) (Cox & Mann, 2008). The derived peak list was searched with the built-in Andromeda search engine (Cox et al., 2011) against the reference *C. neoformans* H99 proteome downloaded from Uniprot (http://www.uniprot.org/) (Aug. 18, 2018; 7,430 sequences). The following parameters were included: trypsin enzyme specificity with a maximum of two missed cleavages, a minimum peptide length of seven amino acids, fixed modifications, including carbamidomethylation of cysteine, and variable modifications, including, methionine oxidation and N-acetylation of proteins and split by taxonomic ID. Peptide spectral matches were filtered using a target-decoy approach at a false-discovery (FDR) of 1% with a minimum of two peptides required for protein identification. Relative label-free quantification (LFQ) and match between runs were enabled; the MaxLFQ algorithm used a minimum ratio count of 1 (Cox et al., 2014). The mass spectrometry proteomics data have been deposited in the PRIDE partner repository for the ProteomeXchange Consortium with the data set identifier: PXD013894.

Statistical analysis of the MaxQuant-processed data was performed using the Perseus software environment (version 1.6.2.2) (Tyanova et al., 2016). Data were prepared by filtering for reverse database matches, contaminants, and proteins only identified by site, followed by log_2_ transformation of LFQ intensities. Filtering for valid values (three of four replicates in at least one group) was performed, and missing values were imputed from the normal distribution (width, 0.3; downshift, 1.8 standard deviations). A Student’s *t*-test was performed to identify proteins with a significant differential expression (*p-value* < 0.05) (S_0_ = 1) between samples employing a 5% permutation-based FDR filter.

### *In vitro* macrophage survival assay

Fungal survival after engulfment by THP-1 (ATCC #TIB-202) cells *in vitro* was assessed as in (Santiago-Tirado et al., 2015) with only minor modifications: First, since the cryptococcal strains were grown in G418 for plasmid maintenance, they were washed extensively with PBS to remove all traces of drug before being opsonized, adjusted to an MOI of 10, and added to the THP-1 cells. Second, for CFU determination, 0.1% Triton-X100 (which had no effect on the cryptococcal cells) was used instead of SDS to lyse the THP-1 cells.

### Infection studies

All animal studies were reviewed and approved by the Animal Studies Committee of Washington University School of Medicine and conducted according to the National Institutes of Health guidelines for housing and care of laboratory animals. Strains to be tested were cultured overnight in YPD medium, collected by centrifugation, washed in PBS, and diluted to 10^6^ cells/ml in PBS for intranasal inoculation (50 µl) into 4-6 week-old female C57BL/6 mice (National Cancer Institute) that had been anesthetized with a combination of ketaset-HCl and xylazine. Initial inocula were plated to confirm CFUs. To assess long-term survival, infected animals were weighed 1 h post-infection and at least every other day afterwards. Mice were sacrificed if their weight fell below 80% of peak (an outcome which in this protocol precedes any signs of disease) or at the end of the study (10 weeks). To measure organ burden, infected mice were monitored as above for 14 days, at which point lungs and brains were harvested from all mice, homogenized in PBS, and serial dilutions of the homogenate plated on YPD agar for enumeration of CFU.

